# A numerical framework for genetic hitchhiking in populations of variable size

**DOI:** 10.1101/2021.03.25.437048

**Authors:** Eric Friedlander, Matthias Steinrücken

## Abstract

Natural selection on beneficial or deleterious alleles results in an increase or decrease, respectively, of their frequency within the population. Due to chromosomal linkage, the dynamics of the selected site affect the genetic variation at nearby neutral loci in a process commonly referred to as genetic hitchhiking. Changes in population size, however, can yield patterns in genomic data that mimic the effects of selection. Accurately modeling these dynamics is thus crucial to understanding how selection and past population size changes impact observed patterns of genetic variation.

Here, we model the evolution of haplotype frequencies with the Wright-Fisher diffusion to study the impact of selection on linked neutral variation. Explicit solutions are not known for the dynamics of this diffusion when selection and recombination act simultaneously. Thus, we present a method for numerically evaluating the Wright-Fisher diffusion dynamics of two linked loci separated by a certain recombination distance when selection is acting. We can account for arbitrary population size histories explicitly using this approach. A key step in the method is to express the moments of the associated transition density, or sampling probabilities, as solutions to ordinary differential equations. Numerically solving these differential equations relies on a novel accurate and numerically efficient technique to estimate higher order moments from lower order moments.

We demonstrate how this numerical framework can be used to quantify the reduction and recovery of genetic diversity around a selected locus over time and elucidate distortions in the site-frequency-spectra of neutral variation linked to loci under selection in various demographic settings. The method can be readily extended to more general modes of selection and applied in likelihood frameworks to detect loci under selection and infer the strength of the selective pressure.

## 1 Introduction

Natural selection acting on an allele alters its frequency in a population over time. Beneficial alleles tend to increase in frequency, whereas deleterious alleles tend to decrease in frequency. More general forms of diploid selection like dominance, recessive, heterozygote advantage, or underdominance yield more intricate behavior. Due to chromosomal linkage, the dynamics of the allele under selection impact the allele frequency trajectories at nearby neutral loci, thus each selection regime results in a characteristic pattern of observed genetic variation. For example, in the case of strong positive selection, the beneficial allele can quickly sweep to fixation. This will cause closely linked neutral sites to increase in frequency as well, despite the alleles being neutral, in a processes referred to as genetic hitchhiking (Maynard Smith and Haigh, 1974; Kaplan et al., 1989). Typically, hitchhiking results in a decrease in genetic diversity around the selected locus. This can be quantified, for example, by examining the heterozygosity around a selected locus, or inspecting the local site-frequency-spectrum (SFS), see for example (Kim and Stephan, 2002, Fig. 2) or (Fay and Wu, 2000, Fig. 1). Numerous successful tools that detect genetic variation under selection in genome-wide sequence data have been implemented to scan for these characteristic patterns, see for example Pavlidis and Alachiotis (2017) or Hejase et al. (2020) for a review.

**Figure 1:**
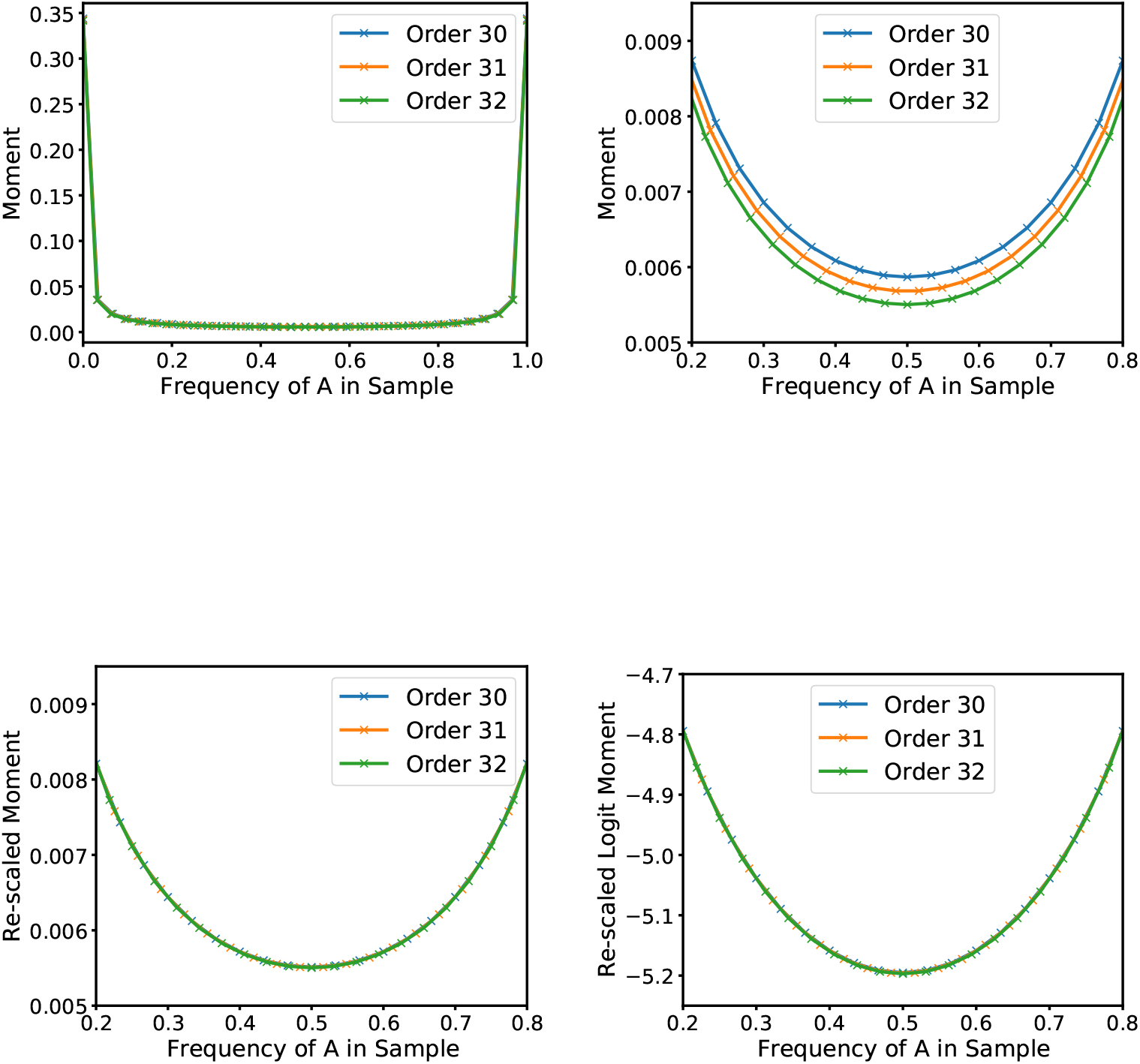
Graphical representation of the *linear-logit* approximation scheme in the single-locus case. Properly rescaled moments of a certain order can be used to approximate moments of higher order.

**Figure 2:**
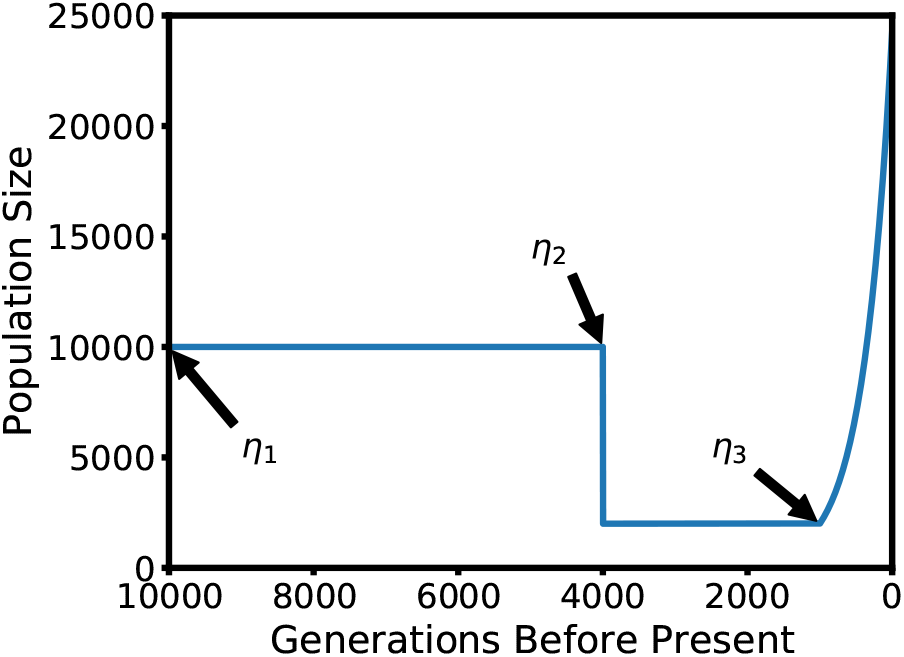
Stylized Out-of-Africa demographic model. The ancestral population size is 10,000, which is reduced to 2,000 during a bottleneck 4,000 generations ago. Exponential growth until the present starts 1,000 generations ago.

While selective sweeps leave very distinct patterns, different scenarios of selection may lead to more subtle signals. For example, balancing selection tends to maintain the selected allele at intermediate frequencies over long evolutionary times (Charlesworth, 2006), whereas selection on polygenic traits is thought to act through concerted small adjustments of frequencies at many loci that affect a trait (Pritchard et al., 2010). In addition to the mode of selection, the demographic history of the population also impacts patterns of genetic diversity and in some cases leads to changes in allele frequencies that mimic the impact of selection (Williamson et al., 2005). Thus, developing methods capable of capturing the dynamics of the interplay of selection, variable population size, and chromosomal linkage is vital to interpreting observed genetic variation and elucidating the underlying population genetic and evolutionary processes.

To this end, we present a numerical method for studying the dynamics of haplotype frequencies at two linked loci in a population of changing size. The loci are separated by a given recombination distance and subject to selective pressure. Our approach can be used to accurately and efficiently compute quantities of interest, including expected allele or haplotype frequency trajectories, and expected heterozygosity. Moreover, it can be used to study statistics of linkage disequilibrium (LD), fundamental to many population genetic tools, for example, to characterize linkage blocks in human genetic data (Berisa and Pickrell, 2015), infer recombination rates (McVean et al., 2002; Spence and Song, 2019), infer complex demographic histories (Ragsdale and Gutenkunst, 2017; Ragsdale and Gravel, 2019), or investigate the polygenic and pleiotropic architecture of complex traits (Bulik-Sullivan et al., 2015; Zabad et al., 2021). Furthermore, our method enables exploring the effects of linked and background selection on the site-frequency-spectrum (SFS), another statistic central to many population genetic analyses aimed at either inferring population genetic parameters from neutral site frequency spectra (Gutenkunst et al., 2009; Kamm et al., 2020) or characterizing selection directly (Boyko et al., 2008; Zivkovic and Stephan, 2011). Many of these and other LD- or SFS-based tools operate under the assumptions of neutrality, and thus it is vital for the correct interpretation of the results to understand how direct and linked selection affects these statistics, while correctly accounting for underlying complex demographic histories. In addition, our methodology provides a tractable way of calibrating existing methods for detecting selection, for example, by determining the size of the genomic footprint for different modes of selection under a given demographic history.

Our method is based on the two-locus Wright-Fisher model, and its diffusion approximation, which has found widespread application in population genetics (see e.g. Ewens, 2010; Durrett, 2008). The Wright-Fisher diffusion is a stochastic process that describes the random dynamics of allele or haplotype frequencies in a population under genetic drift, and can explicitly account for selection, recombination, mutation, and arbitrary population size histories. The dynamics of the Wright-Fisher diffusion can be described by a partial differential operator, or infinitesimal generator. The generator defines a pair of associated partial differential equations (PDEs) known as the Kolmogorov forward and backward equations. The solutions to these equations yield the transition density of the Wright-Fisher diffusion, which in turn gives the likelihoods of allele frequency changes over time. However, no analytic solutions are known to these equations, even in the single-locus case. Instead, numerical approaches have been applied to compute this transition density (Kimura, 1955b,a; Song and Steinriicken, 2012; Bergman et al., 2018; Zhao et al., 2013; He et al., 2020a) or to compute likelihoods of observed genetic data (Jenkins and Song, 2012; Steinrücken et al., 2014; Schraiber et al., 2016; He et al., 2020b). Approaches based on the Wright-Fisher diffusion to study two-locus models when selection is acting have been applied before, for example by Barton and Etheridge (2004) or Stephan et al. (1999), but not in scenarios with variable population size.

To motivate our approach, we note in particular the framework used by Williamson et al. (2005) and Gutenkunst et al. (2009), who, in the single locus case, first solved the PDE using a finite difference scheme to obtain the transition density of the Wright-Fisher diffusion. The authors then integrated the resulting solution numerically to obtain the expected SFS, which in turn was used for likelihood-based inference of demographic parameters and selection coefficients. This approach was later extended by Ragsdale and Gutenkunst (2017) to the two-locus case under neutrality. The primary challenge with this approach, however, is that it can become prohibitively computationally expensive and inaccurate, especially in the two-locus case, due to the high dimensionality required for the solution scheme and accumulation of errors in the subsequent numerical integration. Evans et al. (2007) introduced a moment-based approach that circumvents this two-step procedure. The authors derived a system of ordinary differential equations (ODEs) that can be solved more efficiently to directly compute the moments of the diffusion, which can be used to obtain the expected SFS or the probabilities of obtaining given sample configurations. Jouganous et al. (2017) extended this approach to compute the joint SFS for multiple populations. While their approach can in principle include the action of selection, they focus on the inference of complex demographic histories. One hurdle that arises with the inclusion of selection is that the system of ODEs does not “close.” That is, the dynamics of the moments of a certain order *n* depends on moments of order *n* +1. The authors address this issue through a Jackknife approximation in which the higher-order moments are approximated from those of lower order.

A straightforward extension of this technique to the two-locus case also results in equations that do not close, due to recombination and selection. However, one can consider alternative representations of the moments which close under recombination but not selection. To this end, Ragsdale and Gravel (2019) use an extension of the moments presented by Hill and Robertson (1968) to derive a framework for the estimation of complex demographic histories using multi-population two-locus summary statistics. Kamm et al. (2016) use a related set of moments, derived in a coalescent framework, to perform fine-scale recombination rate estimation. Although these alternative moments do close the system of ODEs under recombination, they also increase its dimension. Since these alternative moments do not close under selection, one would still need to rely on a Jackknife technique to approximate higher order moments to capture the selection dynamics. For the approximation to be accurate, moments of sufficiently high order have to be considered. However, due to the high dimensionality, increasing the order of these alternative moments can become prohibitive, making it computationally expensive to accurately include selection in this approach.

In this work, we introduce a numerical framework to compute the trajectories of the two-locus moments of the haplotype frequencies, or haplotype sampling probabilities, under the Wright-Fisher diffusion subject to genetic drift, mutation, recombination, and selection. This method can simultaneously capture the impact of selection, recombination, and changing population size on higher order moments, or sampling probabilities, making it a valuable tool for studying patterns of genetic variation arising in populations under selection. To circumvent the high dimensionality required for alternative moment representations, we instead use the regular moments (defined below) in our implementation, combined with a novel efficient approximation of higher order moments to “close” the dynamics for recombination and selection. A similar approach, including selection, is outlined in Section S1.3 of the Appendix in Ragsdale and Gravel (2019). Here, we give an alternative derivation which relies on Dynkin’s formula. Moreover, Ragsdale and Gravel (2019) focus on the neutral case in their study, but the approach has recently been used by Ragsdale (2021) to explore how epistasis and dominance shape expected patterns of signed linkage disequilibrium in protein coding variants. In this work, however, we explore the dynamics of one selected locus and its effect on linked neutral variation in populations with complex demographic histories. Specifically, we investigate a scenario with a population bottleneck followed by exponential population growth, relevant to the Out-of-Africa bottleneck and subsequent population expansion in human evolution. By comparing to simulations, we show that our method can accurately and efficiently compute expected allele and haplotype frequency trajectories, heterozygosity, linkage disequilibrium, and the local SFS in a window around a selected locus. Thus, the outlined method provides the means to further our understanding of patterns observed in genomic data from populations subject to selective pressure, in which the history of changing population sizes may obscure or confound the signal of selection.

This paper is organized as follows. In Section 2.1, we derive the system of ODEs for the moments of the two-locus diffusion by applying Dynkin’s formula to the infinitesimal generator of the Wright-Fisher diffusion. We present our novel moment closure technique in Section 2.2, and demonstrate its accuracy by comparing with several alternative approaches in Section 3.1.1. In Section 3.1, we show that our method accurately computes the expected trajectories of allele frequencies, haplotype frequencies, heterozygosity and linkage disequilibrium across a broad range of parameters by comparing it to simulated trajectories. In addition, in Section 3.2, we show that the method accurately captures the impact of selection at a given locus on the local SFS in a window around the selected site. Finally, in Section 4 we discuss possible applications of our method in composite likelihood frameworks to detect and infer the strength of selection, as well as other strategies for how our method can improve existing inference techniques.

## 2 Methods

### 2.1 Moment Equations

In this section we first give a description of the Wright-Fisher diffusion, which is the mathematical framework we use to develop our method. This is followed by a derivation of the system of ODEs for the moment dynamics, as well as a description of the moment-closure technique essential to the method.

#### 2.1.1 Wright-Fisher Diffusion

We first introduce the two-locus Wright-Fisher diffusion. Consider a panmictic, diploid population of size *N_t_* (2*N_t_* gametes), at time *t* ≥ 0, where time is measured in 2*N*_0_ generations, where *N*_0_ is the size of the reference population. We model the dynamics of two linked loci, locus *A* and locus *B*. Let *E^A^* = {1,…, *K_A_*} and *E^B^* = {1,…, *K_B_* } represent the set of alleles at locus *A* and locus *B*, respectively. It follows that the set of possible two-locus haplotypes is given by *E* = *E^A^* × *E^B^*, with *K* = |*E*|. Any haplotype *i* ∈ *E* can be represented by the 2-tuple (*i_A_, i_B_*), where *i_A_* and *i_B_* are the alleles at locus *A* and *B*, respectively. Let ***X***(*t*) = {*X_k_*(*t*)}_*k*∈*E*_ be the vector of frequencies of the haplotypes in the population at time *t*, where ***X***(*t*) takes values in the (*K* – 1)-dimensional simplex,

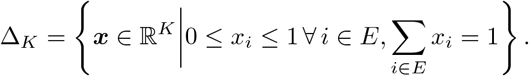

For moderate to large population sizes, the dynamics of **X**(*t*) can be modeled using the Wright-Fisher diffusion, see for example (Durrett, 2008, Ch. 7) or (Ewens, 2010, Ch. 4). Let 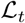 be the infinitesimal generator of the Wright-Fisher diffusion, which we define in more detail below. In general, this generator depends on the time *t*, allowing us to model changing population sizes. We refer the reader to (Karlin and Taylor, 1981, Ch. 15.2) or (Ethier and Kurtz, 2009, Ch. 10) for a more detailed introduction on the concept of infinitesimal generators for diffusions. The transition density of the Wright-Fisher diffusion, 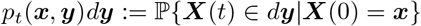 for ***x, y*** ∈ Δ_*K*_, can be obtained as the solution to the corresponding backward equation,

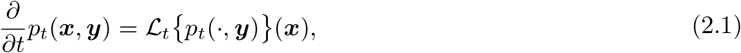

for **y** fixed and 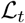 acting on *p_t_*(***x**, **y***) as a function of ***x***. Alternatively, the density can be characterized through the corresponding forward equation where ***x*** is fixed and the adjoint of 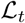 is applied to *p_t_*(***x**, **y***) as a function of ***y***.

In this work, we model the dynamics in a population subject to genetic drift, mutation, natural selection, and recombination. The Wright-Fisher diffusion has the property that the infinitesimal-generator is linear in these population genetic processes. Namely, the generator can be decomposed as follows,

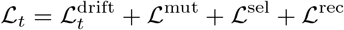

where 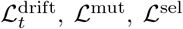, and 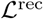 represent the contribution to the generator from genetic drift, mutation, selection, and recombination, respectively. These four operators are given as follows:

##### Genetic Drift Operator

Setting 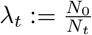, the operator representing genetic drift is

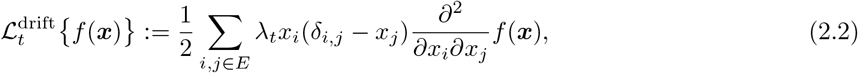

where *δ_i,j_* is the Kronecker delta, see for example (Durrett, 2008, Ch. 8.1). Here the dependence of *λ_t_* on time *t* allows the model to capture changing population size by varying the strength of genetic drift.

##### Mutation Operator

For a model of recurrent mutation, let *m_i,j_* for *i,j* ∈ *E* be the per generation probability that a gamete carrying haplotype *i* mutates to haplotype *j*. Let 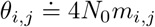 be the population scaled mutation rate from haplotype *i* to *j* and let 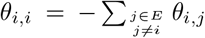. The operator corresponding to mutation can then be written as,

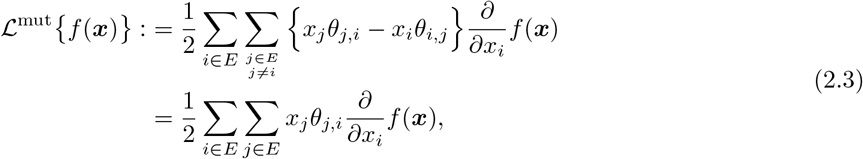

see for example (Durrett, 2008, Ch. 8.1). This formulation allows a very general mutation model, where in principle, the alleles at both loci could mutate at the same time. If mutation rates are low, and only one locus mutates at a time, we could use the special case *θ_i,j_* = *θ_i_A_,j_A__*, if *i_B_* = *j_B_* and *θ_i,j_* = *θ_i_B_,j_B__*, if *i_A_* = *j_A_*.

Unfortunately, the generator for models with non-recurrent mutation in the limit of an infinite population size is often not well-defined in the literature, but see Evans et al. (2007). This is especially true when considering the variation at a particular pair of loci as we do here. We nonetheless introduce the correct implementation of a non-recurrent mutation model at two linked loci in our numerical framework in Appendix A.2.2 using arguments related to the diffusion, but slightly different from the other cases considered here. Note that this implementation of the non-recurrent mutation model results in a non-linear ODE. However, for convenience, we will use the notation of a linear ODE in the remainder.

##### Selection Operator

We assume a model of general diploid selection. To this end, let *s_i,j_* for *i,j* ∈ *E* be the fitness advantage of a diploid individual carrying the haplotypes *i* and *j*, that is, the probability that this individual is the parent of an offspring individual in the next generation is proportional to 1 + *S_i,j_*. Furthermore, let *σ_i,j_* = 4*N*_0_*s_i,j_* be the population-scaled relative fitness of an individual with haplotypes (*i,j*). Without loss of generality, we can set *s_κ,κ_* = 0 = *σ_κ,κ_*. Define the marginal fitness of haplotype *i* ∈ *E* in a population with haplotype frequencies ***x*** as,

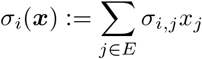

and the mean fitness of the population 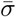 as,

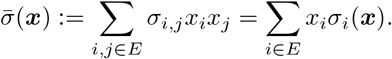

Using this notation, the selection operator can be expressed as,

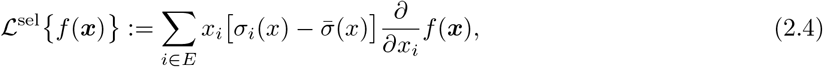

see for example (Steinrücken et al., 2013).

##### Recombination Operator

Let *r* be the per generation probability of recombination between locus *A* and *B*. For allele *i_A_* ∈ *E^A^* at locus *A* and allele *i_B_* ∈ *E^B^* at locus *B*, we define the marginal allele frequencies as,

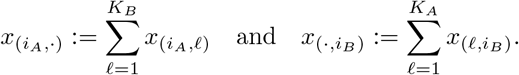

The recombination operator is then given by,

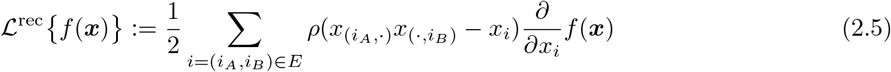

where 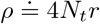 is the population scaled recombination rate between *A* and *B*, see (Jenkins and Song, 2012).

#### 2.1.2 Dynkin’s Formula and Moment Dynamics

The transition density *p_t_*(***x**, **y***) of the Wright-Fisher diffusion describes the dynamics of the haplotype frequencies at two linked loci. Ideally, one would obtain *p_t_*(·, ·) analytically by solving the Kolmogorov forward or backward equations, however, no analytical solution is known for these PDEs in general. Numerically solving equation (2.1), for example by using finite difference methods, is computationally expensive and challenging in many high-dimensional settings (Gutenkunst et al., 2009; Ragsdale and Gutenkunst, 2017). In order to circumvent solving the PDE of the forward equation (2.1), an alternative technique presented by Evans et al. (2007) provides a less computationally intensive method of directly computing the moments of the transition density. These moments also yield the probability of obtaining a given sample of haplotypes from the population, and thus are also referred to as sampling probabilities. Conveniently, in many applications these moments are the desired quantities, and computing the transition density is just an intermediate step to obtain them (Gutenkunst et al., 2009; Ragsdale and Gutenkunst, 2017; Williamson et al., 2005). The moments can also be used to compute the expected values of many statistics obtained from population genetic data, or for likelihood-based inference of parameters.

Suppose that at time *t* ≥ 0 one obtains a sample of haplotypes **n** = {*n_i_*}_*i*∈*E*_ from the population, where *n_i_* denotes the number of haplotypes of type *i* ∈ *E* in the sample. Setting 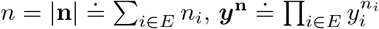, and denoting by ***y***_0_ the haplotype frequencies at time 0, the probability of obtaining such a sample is given by,

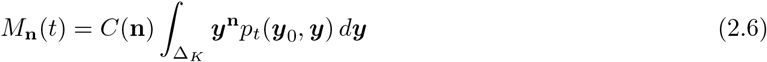

where 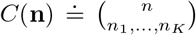 is the corresponding multinomial coefficient. We will frequently refer to *M*_n_ as defined above as a “moment of order *n*” and find it helpful to define the vector of all moments of order *n* as follows. For each 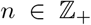, define the set 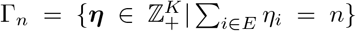, and define the vector **M**_*n*_ = {*M_n_*(*t*)}_*n*∈Γ_n__ as the collection of all moments of order *n*. The evolution of **M**_*n*_(*t*) is given as the solution to a system of ODEs derived from the Wright-Fisher diffusion. In previous works, these ODEs have been derived using integration-by-parts (Evans et al., 2007) or the transition probabilities in a finite population (Jouganous et al., 2017; Ragsdale and Gravel, 2019). We give an alternate, more convenient, derivation using Dynkin’s formula, see for example (Karlin and Taylor, 1981, Ch. 15) or (Øksendal, 2003, Lemma 7.3.2).

##### Theorem 2.1

(Dynkin’s Formula). *Let **X** be a diffusion with generator 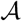 and let f be a compactly-supported and twice differentiable function with continuous second derivative. Then if X*(0) = *x*,

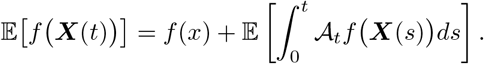

To obtain the requisite system of ODEs describing the evolution of **M**_*n*_(*t*), we apply Dynkin’s formula to functions *f* of the form of the integrand in equation (2.6):

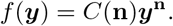

This yields,

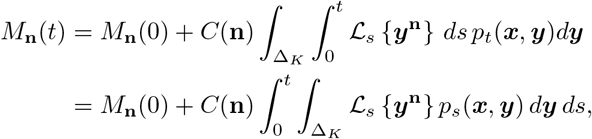

where the second equality follows from Fubini’s Theorem. Taking the derivative with respect to time *t* on both sides then gives,

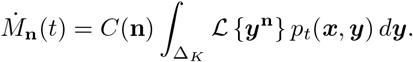

The following theorem is the key representation used in our numerical method.

##### Theorem 2.2.

*The derivative of the moment-vector of order n*, 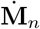, can be written as a linear combination of moments of order *n*, *n* + 1, *and n* + 2 *in the form*,

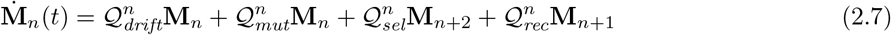

*where* 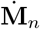 *is the vector of derivatives for each entry of* **M**_*n*_. *Furthermore, when only genic selection is considered* 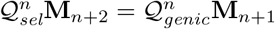 *and the order n* + 2 *moments are no longer needed*.

The proof of this theorem relies on carefully rearranging terms for each individual moment 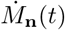 and aggregating across moments (see Appendix A for details) to show that,

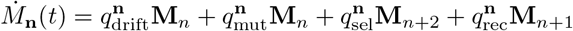

where each *q*^n^ is a row vector of appropriate dimension corresponding to the four population genetic processes of interest in this work. The row vectors *q*^n^ are then aggregated into matrices 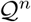. A detailed proof is provided in Appendix A. With this representation, we can obtain the moments of the transition density by solving this system of ODEs, which is numerically more tractable than solving the multi-dimensional PDE of the Wright-Fisher diffusion. However, note that moments of order higher than *n* appear on the right-hand side of equation (2.7). The system of ODEs does not “close” in that the derivative in *t* of the moments of order n depends on moments of order higher than *n*. The lack of closure is due to the action of selection and recombination. Therefore, in order to solve these ODEs, one would need to know all moments of all orders. We detail a novel alternative solution to this problem in Section 2.2, inspired by the approach of Ragsdale and Gravel (2019), in which higher order moments are estimated from those of lower order. Using this approximation effectively “closes” the ODEs and thus makes it tractable to numerically solve the ODEs for a given order *n*.

### 2.2 Moment Closure

In this section, we detail our novel approach to approximate higher order moments and close the ODEs described in the previous section. As we will compare our moment-closure approach to others, it will be convenient to refer to this method as *logit-linear* throughout the paper. Our approach is motivated by the fact that properly rescaled moments converge to the underlying distribution. The following convergence lemma makes this statement more precise in the one-dimensional case:

#### Lemma 2.3.

*Let μ be a probability measure on* [0, 1] *that has a density f*: 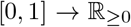. *For the sequence of approximating measures,*

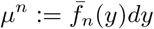

*with density*,

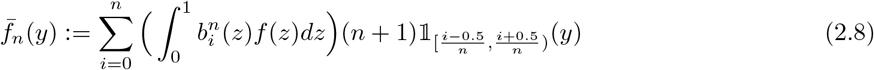

*where,*

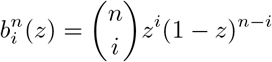

*are the Bernstein polynomials, we have that*

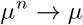

*converges weakly as n* → ∞.

Here 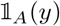 is the indicator function which is equal to 1 if *y* ∈ *A*, and equal to 0 if *y* ∉ *A*. We provide a proof of this lemma in Supplementary Material S1. Note that the factor *n* +1 in equation (2.8) compensates for the fact that the individual moments computed by the integral converge to 0. This factor does not need to be exactly *n* + 1, every factor that grows at same rate as *n* would lead to the same results. We choose *n* + 1 for convenience.

To illustrate Lemma 2.3 in practice, consider a single locus with two alleles *A* and *a*. Suppose that the population frequency of the *A* allele follows a Beta distribution with parameters *α* = *β* = 0.1. The Beta distribution is the stationary distribution under the mutation model considered here. Our aim is to approximate moments of order *n* + *m* from those of order *n*. To illustrate this, we depict in Figure 1(a) the vectors of moments of order 30, 31, and 32. The *x*-coordinate for the *i*-th moment of order *n* is 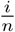, and we interpolate linearly between these values. Since the moments also represent probabilities of obtaining a certain sample configuration, the fraction 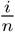 is also the frequency of the *A* allele in the sample. Note that in Lemma 2.3, the density 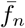 is defined as a piece-wise constant function, whereas we use linear interpolation here. However, the result from Lemma 2.3 would hold for any suitable interpolation scheme, so we proceed with the linear interpolation for illustration purposes.

We observe that the shapes of the curves for different *n* are similar, and the values are reasonably close to each other. However, as can be seen in Figure 1(b), when focusing on the values for intermediate frequencies, the exact values indeed differ and decrease as the order *n* increases. As Lemma 2.3 suggests, the discrepancy between moments of different orders can be compensated for by rescaling the moments appropriately. Indeed, Figure 1(c) shows the same curves, but the moments of order *n* = 30 and *n* + 1 = 31 are multiplied by 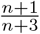 and 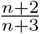, respectively. The resulting curves align well. However, note that the moments are small, and will further decrease with increasing *n*. Moreover, the moments also represent probabilities and the values are in the interval [0,1]. Thus, we can apply the `logit` transformation

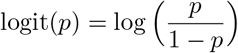

to improve the resolution for small values and further improve the fit. Figure 1(d) shows the resulting transformed values, and suggests approximating higher order moments by interpolating the `logit`-transformed rescaled values.

For the two-locus case, we require a multi-dimensional version of Lemma 2.3:

#### Conjecture 2.4.

*Let μ be a probability measure on* Δ*_κ_ with density f*: 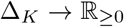. *Define the measures μ_n_ on* Δ*_κ_ with densities*

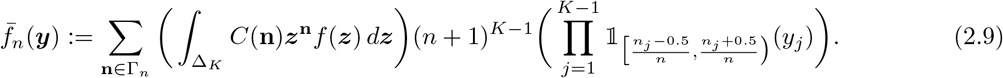

*Then*

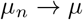

*converges weakly*.

This conjecture extends the results of Lemma 2.3 to multiple dimensions, but we do not provide an explicit proof for it here.

By substituting the transition density of the Wight-Fisher diffusion *p_t_*(***x**, **y***) for the density *f*(***y***) in equation (2.9), we observe that when the moments of the Wright-Fisher diffusion are properly rescaled they approximate its transition density as the moment order increases. Again, the factor (*n* + 1)^*K*-1^ reflects the fact that individual moments converge to 0, since the total probability mass is spread out over an increasing number of configurations as *n* increases. The factor compensates for this effect.

We use Conjecture 2.4 to motivate our moment approximation technique as follows. Convergence of the rescaled moments to the transition density implies that the elements of the approximating sequence are increasingly close to the limit. The individual elements of this sequence of rescaled moments can be used as approximations to each other. Thus we can accurately approximate *M*_n_′(*t*) for a given configuration **n**′ with |**n**’| = *n* + 1 by interpolating the values *M*_n_(*t*) for certain configurations **n** with |**n**| = *n* “close” to **n**′. Here, we use linear interpolation of the `logit` of the values of *M*_n_(*t*). We now explicitly define our *logit-linear* approximation in the general case.

#### Logit-linear Interpolation

To define our moment approximation scheme in the two-locus case, we restrict ourselves to the setting in which both loci are bi-allelic for convenience, that is |*E^A^*| = |*E^B^*| = 2 and *K* = 4. Then, the moment vectors can be represented as three dimensional vectors in the 3-simplex Δ_4_. As suggested by Conjecture 2.4 and outlined in the single-locus example, we will (1) rescale the moments so that the magnitude is comparable across orders, (2) apply the `logit` transformation to the rescaled moments, (3) perform linear interpolation, and (4) invert the `logit` transformation to obtain the approximate moments of higher order. For numerical stability, we will also (5) rescale the approximate moments to sum to 1.

Suppose we have moments of order *n*, call them **M**_n_ as above, suppressing the dependence on time, and we wish to estimate moments of order *n* + *m*, **M**_*n+m*_. In the case of genic selection and recombination (analyzed in Section 3.1) we will only need to increase the moment order by 1, that is *m* = 1, while for general models of selection *m* = 2 is needed. In order to rescale **M**_*n*_ to a comparable magnitude as **M**_*n+m*_ we will multiply **M**_*n*_ by,

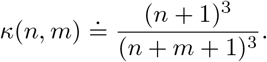

Focusing on the details of the interpolation step, recall that the moment vectors **M**_*n*_ and **M**_*n+m*_ are indexed by the sets Γ_*n*_ and Γ_*n+m*_, respectively. For any 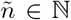, define sets 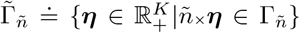. Namely, these are the same as the sets Γ_*n*_ and Γ_*n+m*_ with the elements rescaled so that they are elements of Δ_4_, the 3-dimensional simplex. Whereas Γ_*n*_ is the set of all moment powers that add up to *n*, 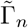 represents the corresponding configurations in the frequency space Δ_4_. For every 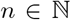 let 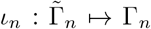 map the elements in 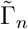 to the corresponding configuration in Γ_*n*_. We call this the “sample frequency representation” of a moment since it represents the frequency of each haplotype in a sample of size *n*. Define the function *f_n,m_*: 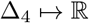 where for every 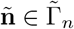,

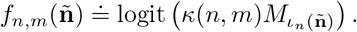

Specifically *f_n,m_* maps the scaled indices 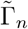 to their corresponding elements of **M**_*n*_, rescaled by *κ*(*n, m*) and with the `logit` transformation applied. Given ***l*** ∈ Γ_*n+m*_, *M_**l**_* is estimated by first computing the linear interpolation of 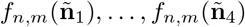, where 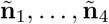 are the four elements of 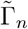 closest to 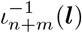, and then applying the inverse of the `logit` transformation. We perform this process to obtain each element of **M**_*n+m*_. The final step is to re-normalize the estimated moments **M**_*n+m*_ to sum to one. We found that this scheme increased accuracy and stability when numerically solving the moment ODEs. In Section 3.1.1, we will provide comparison to the following alternative estimation techniques.

##### 2.2.1 Alternative Techniques

In this section we give a brief outline of other moment approximation techniques that we considered. While these methods all provide viable means of estimating higher order moments, the *logit-linear* method showed the best performance, as we will demonstrate in Section 3.1.1.

###### Linear Interpolation (no logit)

The first alternative considered, referred to as *linear*, is the same as *logit-linear* without applying the `logit` transformation. Specifically, the steps for linear are (1) rescale the moments of order *n*, (2) perform linear interpolation on the rescaled moments, (3) rescale the resulting approximate moments of order *n* + *m* to sum to 1.

###### “Jackknife” of Jouganous et al. (2017), Ragsdale and Gravel (2019)

This moment closure technique was first introduced for the single-locus setting by Jouganous et al. (2017) following ideas presented by Gravel et. al. (2014). The basic approach is to approximate the distribution of the population allele frequency underlying the moments using a quadratic function *a*_0_ + *a*_1_*x* + *a*_2_*x*^2^ with coefficients ***a*** = [*a*_0_, *a*_1_, *a*_2_], which results in expressions for the moments of any order in terms of **a**. A given moment of order *n* + *m*, call it *m_i_*, can be approximated as follows: Select the three moments of order *n* which are “closest” (in the sense of their sample frequency representation), say 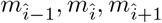. Since these moments of order n are known quantities, one can derive a system of equations and solve for the parameters **a**. In turn, these parameters can then be used to compute *m_i_*. While assuming the distribution is quadratic may seem restrictive, note that since the **a**’s are not fixed a priori and are recomputed for each *m_i_*, this amounts to assuming the distribution is only locally quadratic, a much milder assumption. For more information we refer to reader to Jouganous et al. (2017).

In Appendix S1.3 of (Ragsdale and Gravel, 2019), the authors present a two-locus implementation of this scheme. Assuming bi-allelic loci, the quadratic function for the population allele frequency is then a three dimensional function on Δ_4_ and requires 10 coefficients. Thus, in order to estimate a moment of order *n* + *m* (indexed by (*i, j, k*)), one must select the 10 closest moments of order *n* (indexed by 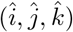). The authors further specify a constraint stating that three separate values for each coordinate (i.e. 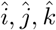) must be included in the set used to estimate the quadratic coefficients. We will refer to this method as *jackknife* and refer the reader to (Ragsdale and Gravel, 2019, Appendix S1.3.5) for a more detailed explanation. In addition, we consider the estimator without the added constraint that each coordinate takes three values which we will call *jackknife-unconstrained*.

###### Least Squares Projection

Lastly, we also considered a method that we refer to as *least-squares*. To this end, we say that **M**_*n+m*_ and **M**_*n*_ are “parsimonious” if **M**_*n*_ is equal to the moments of order *n* computed from the larger samples in **M**_*n+m*_. Note that if we view the moments as sampling probabilities then **M**_*n*_ can be computed by downsampling from **M**_*n+m*_. In this case the two vectors will satisfy a set of identities relating them to one another, see Supplementary Material S2.2 for a more detailed description. One can then define a matrix 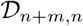 representing these relations such that 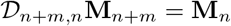 holds if they are parsimonious.

In the *least-squares* estimator we select the vector of moments of order *m* + *n* which is parsimonious with the moments of order *n* and has the smallest sum of squares. In particular we take the estimate of **M**_*m+n*_ to be the solution to the following optimization problem:

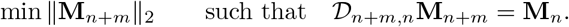

This is a common optimization problem for which closed form solutions and efficient solvers exist. We refer the reader to (Boyd and Vandenberghe, 2018, Ch. 16) for a detailed introduction.

## 3 Results

### 3.1 Simulation studies

In this section, to verify the efficiency and accuracy of our novel method, we compare moments obtained using our numerical solutions of the ODEs with the same quantities obtained from simulations conducted using `SimuPOP` (Peng and Kimmel, 2005; Peng and Amos, 2008). Details on our numerical implementation of a step-wise scheme to solve the ODEs can be found in Supplementary Material S2.1. We note that in order to make the simulations computationally feasible, we follow a standard approach to decrease the population size by a factor of 10 and rescaled all other parameters accordingly. Section 3.1.1 is devoted to validating the model when the genetic variation at the pair of loci separated by a given recombination distance is initialized with fixed haplotype frequencies. We also demonstrate that our novel moment approximation technique outperforms the other candidates outlined in Section 2.2.1 in the simulated scenarios. In Section 3.1.2, we investigate scenarios where the genetic variation at the neutral sites is initialized from the stationary distribution, and vary the recombination distance to exhibit linkage effects as they change along the genome. In both sections we consider the case in which all loci are bi-allelic (i.e. four haplotypes at a pair of loci) and only consider the case of genic selection. We consider a recurrent mutation model in which every allele mutates at rate *m*, that is haplotypes which share the same allele at one locus but differ at the other mutate into each other at rate *m*. We expect similar accuracy if the mutation probabilities would be locus-specific. The demographic model considered here is an idealized approximation of a European population in the Out-of-Africa model (e.g. Jouganous et al., 2017; Gutenkunst et al., 2009). The ancestral population size is 10,000, which is reduced to 2,000 during a bottleneck that starts 4,000 generations before present. The population stays at this reduced size, until 1, 000 generations before present, when it experiences exponential growth at rate .25% per generation until the present, see Figure 2 for a schematic.

By considering different initial times (labeled *η_1_, η_2_*, and *η_3_*) for the introduction of the beneficial allele (10,000 generations before present, at the beginning of the bottleneck, and at the beginning of the growth phase, respectively), and exhibiting the moments at several points in time afterwards, this model essentially allows us to investigate the dynamics of relaxation to stationarity, during the bottleneck only, or just exponential growth, in a single model, thus covering a wide range of demographic scenarios, providing confidence that our method can accurately account for arbitrary population size histories. In all cases we used moments of order 31 for the numerical solutions. This choice resulted from balancing a high moment order for the accuracy of the moment approximation with computational limitations due to the dimension of the moment vector **M**_*n*_ increasing cubically with the moment order.

#### 3.1.1 Fixed Initial Frequencies

In this section, we consider the setting of two loci separated by a recombination distance *ρ* = 4*N*_0_*r*. We consider a scenario with two alleles (*A* and *a*) at the first locus and two (*B* and *b*) at the second locus. At each locus, we use the recurrent mutation model, where an allele mutates into the other with rate *θ* = 4*N*_0_*m* and we consider genic selection on allele *A* with population scaled selection coefficient *σ* = 4*N*_0_*s*, where *s* ranges from 0 to 0.005. In order to validate our results we compare moments obtained from the numerical solution of the ODEs to moments obtained from simulations using `SimuPOP` (Peng and Kimmel, 2005; Peng and Amos, 2008). We consider a variety of different parameter values for the initial haplotype frequencies, selection coefficient, and recombination rate. For the initial haplotype frequencies we vary the initial frequency *x_A_* of the selected allele *A* and initialize the population such that frequencies of haplotypes *AB, Ab, aB*, and *ab* are *x_A_*, 0, 0.5 – *x_A_*, and 0.5, respectively. The different parameter values used are provided in Table 1 and all combinations of parameters values were tested. We chose these to roughly mirror human genetic parameters, with a per generation per locus mutation probability of *m* = 1.25 o 10^-8^, and a per generation recombination probability of *r* = 10^-8^ between two neighboring nucleotide sites. The recombination probability of *r* = 10^-2^ thus roughly corresponds to a physical distance of 1,000,000 bp. In addition, throughout this paper all population scaled parameters (i.e. *ρ, σ, μ*) are given with respect to a reference population size of *N*_0_ = 10, 000, regardless of the details of the demographic model.

**Table 1:** Parameters values that are used to compare the numerical method against simulations.

|  |  |
| --- | --- |
| Initial Frequency of Beneficial Allele ($x_A$) | 0.01, 0.03, 0.05, 0.1 |
| Selection Coefficient ($\sigma$) | 0, 0.01, 0.1, 1, 10, 100 |
| Recombination Rate ($\rho$) | 0.0004 ($r = 10^{-8}$), 400 ($r = 10^{-2}$) |
| Mutation Rate ($\theta$) | 0.0005 ($m = 1.25 \cdot 10^{-8}$) |

##### Comparison of Moment Approximation Methodology

We now compare the moments obtained from numerically solving the ODEs when utilizing different moment approximation techniques to the same moments obtained from the simulations described above. As discussed in Section 2.2, the moment approximation techniques we consider are *logit-linear, linear, jackknife, jackknife-unconstrained*, and *least-squares*. In addition to these methods we test two additional modifications to each, which can be employed in the numerical solution of the ODEs. The first is whether or not to re-normalize the vector of moments after each step of the ODE solver. Specifically, after the solver takes a step the error incurred from the moment approximation may cause the moment vector not to sum to 1, however, we can re-normalize the state vector to sum to 1. The second option is to force the approximated higher-order moment to be “parsimonious” with the lower-order moment. We will defer a detailed description of this modification to Supplementary Material S2.2, but it amounts to projecting the original estimate of **M**_*n*+1_(*t*) onto the space of order *n* +1 moments which reduce to the original **M**_*n*_(*t*) when the order n moments are computed. Note that by definition the *least-squares* approximation technique already satisfies this requirement so the parsimonious projection does not modify this approximation.

The solutions of the ODEs and simulations were compared as follows. For each combination of parameters in Table 1, we performed 1,000 repetitions using `SimuPOP` At the end of each repetition, the expression under the integral in equation (2.6) were computed based on the haplotype frequencies in each replicate. These were then averaged over the 1,000 repetitions to get estimates for all moments of order 31. Furthermore, the respective moments were also computed using the numerical approach to solve the ODE with different moment approximation schemes. In order to evaluate the accuracy of the different approximation schemes, we computed (1) the total absolute error and (2) the mean-squared error (MSE) between the simulated moments and those obtained from numerically solving the respective ODEs. In Table 2 and Table 3, for *logit-linear, linear, jackknife, jackknife-unconstrained*, and *least-squares*, we present the combination of renormalization and parsimonious projection which leads to the best results. In all cases, except for *linear*, the combination of options with the lowest absolute error and lowest MSE were the same. For the case of *linear*, including the parsimonious projection without renomalizing resulted in the best absolute error while excluding parsimonious projection and renormalizing resulted in the best MSE. In Table 2 we report the results where we average the errors across all parameter configurations. We note that *logit-linear* outperforms the other methods on both Absolute Error and Mean-Squared Error, by several orders of magnitude in some cases. Similarly, in Table 3 we report the results of the worst performance across all parameter configurations. Specifically, for each combination of approximation type, renormalization, and parsimonious projection we compute the highest Absolute Error and Mean-Squared Error between the different parameter configurations. We then select the approximation type with the minimum (i.e. the mini-max) of these quantities across renormalization, and parsimonious projection. For this metric, the combination of options with the lowest absolute error and lowest MSE were the same with the exception of *Jackknife-Unconstrained*. Again, the scheme *logit-linear* with renormalization outperforms all other approximation techniques, in some cases by several orders of magnitude. Based on these simulations we chose *logit-linear* as the moment approximation method used in the remainder of this manuscript.

**Table 2:**
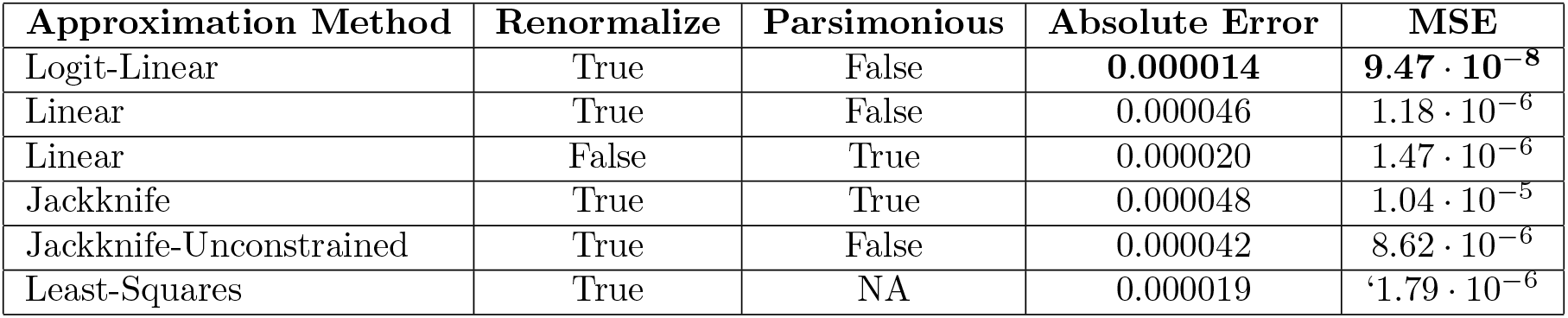
Absolute error and mean squared error comparing simulated moments of order 31 with numerical solutions using different moment approximation schemes, averaged over the parameters given in Table 1.

**Table 3:** Mini-max Absolute error and mean squared error comparing simulated moments of order 31 with numerical solutions using different moment approximation schemes, among the parameters given in Table 1.

| Approximation Method | Renormalize | Parsimonious | Absolute Error | MSE |
| --- | --- | --- | --- | --- |
| Logit-Linear | True | False | **0.000027** | **$4.24 \cdot 10^{-7}$** |
| Linear | True | False | 0.000122 | $8.88 \cdot 10^{-6}$ |
| Jackknife | True | True | 0.000254 | $1.07 \cdot 10^{-4}$ |
| Jackknife-Unconstrained | True | False | 0.000334 | $2.92 \cdot 10^{-5}$ |
| Jackknife-Unconstrained | True | True | 0.000240 | $1.01 \cdot 10^{-4}$ |
| Least-Squares | True | NA | 0.000201 | $6.75 \cdot 10^{-5}$ |

##### Accuracy and Efficiency with Varying Moment Order

To better understand how the choice of the moment order affects accuracy and efficiency of our method, we performed the following analysis. We computed the numerical solutions for different moment orders using our method for all parameter combinations from Table 1. For each moment order, we then down-sampled the vector of moments to a vector of order 10, and computed the error between this vector and the same vector obtained from 1,000 replicated simulations using `SimuPOP`. This analysis corresponds to a situation, where a user would ultimately only be interested in moments of order 10, but the actual order used in the numerical solution is increased to capture the dynamics for recombination and selection more accurately. In Figure 3(a), we depict the error of the moments obtained from the numerical solution for different orders. As expected, the accuracy of the numerical solution of the ODE increases with increasing order, but for larger orders, the gain diminishes. In Figure 3(b), we show the corresponding runtimes, obtained on a 2.0 GHz Intel Haswell computing core. The runtimes increase with increasing moment order. This is expected, as the size of the moment vector increases exponentially with moment order, although the runtimes appear to increase sub-exponentially. Ultimately, the choice of order is a balance between accuracy and efficiency. For the remainder of the paper, we chose *n* = 31 to have a high degree of accuracy, while still computing the solutions in a reasonable amount of time.

**Figure 3:**
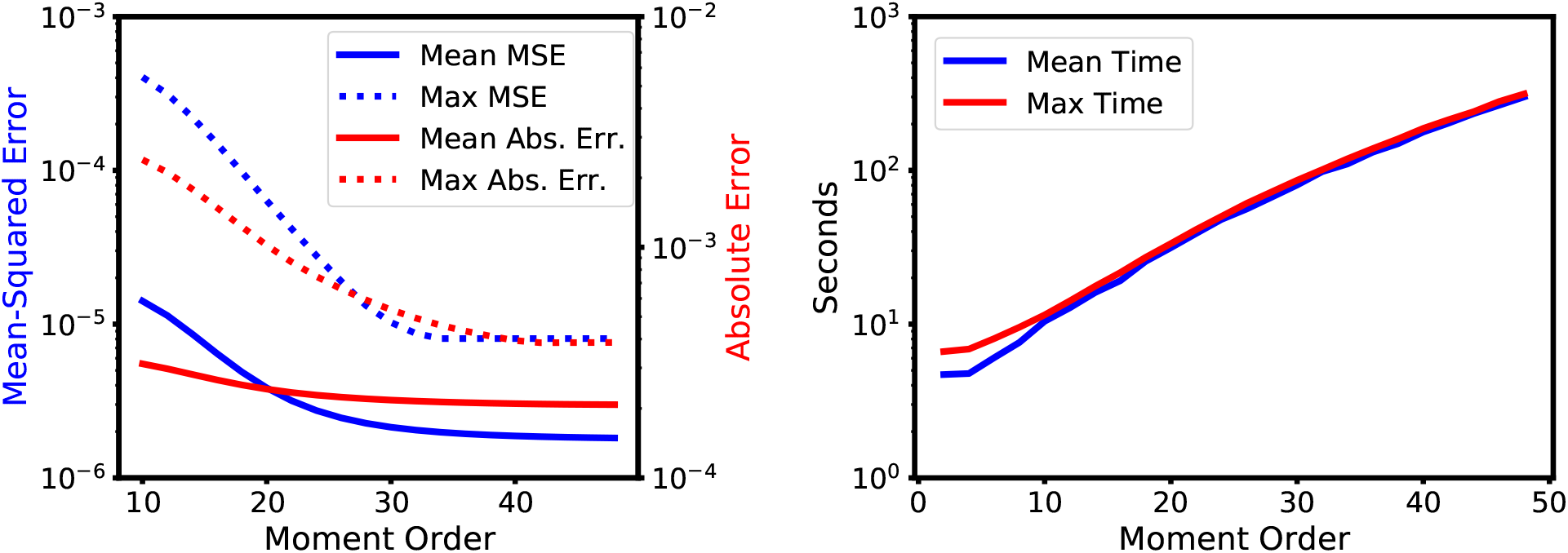
Error and runtime of the numerical solution scheme. The accuracy increases with increasing order, but the runtime also increases. Thus, the choice of the order depends on the user’s preferences regarding this balance.

##### Comparing Trajectories of Moments

In this section, we will present comparisons of trajectories of the moments between the simulations and the numerical solutions of the ODEs. We begin by noting that moments of lower order can be computed from moments of higher order, and can then subsequently be used to compute quantities of interest, for example expected allele frequencies, expected haplotype frequencies, expected heterozygosity or expected linkage disequilibrium. Thus, we compute the numerical solutions of order 31 first, to ensure sufficient accuracy of the moment approximation, and then compute the moments of lower order that we investigate. In Figures 4, 5, 6, and 7, we compare the dynamics of the solutions to the ODEs with simulations starting the dynamics 10,000 generations before present (*η*_1_), 4,000 generations before present (beginning of the bottleneck, *η*_2_), and 1,000 generations before present (beginning of the exponential growth, *η*_3_). We show the expected values computed from the numerical solution of the ODEs and the mean over the 1,000 simulated replicates, also indicating the region of ±1.96 × standard error, corresponding to a 95% confidence interval. Figure 4 displays the expected trajectory of the selected allele *A* computed using the numerical solutions of the ODEs and from the average of the `SimuPOP` simulations. We show the trajectories for *x_A_* = 0.05 and the three largest selection coefficients *σ* =1,10, and 100. We can see that the solution to the ODEs accurately tracks the values from the simulations, and thus captures the population dynamics well. As expected, a larger selection coefficient leads to a faster increase of the allele frequency towards fixation at 1. The increase is slower during the bottleneck (red trajectories), since the efficacy of selection is reduced in the smaller population. In the single-locus case, we can also compute this expected allele frequency more accurately using a different technique developed by Steinrücken et al. (2014). However, this technique can only be applied in the case of constant population size, but we provide comparisons to the numerical solution of the ODE and simulations in Supplementary Material S3.1.2.

**Figure 4:**
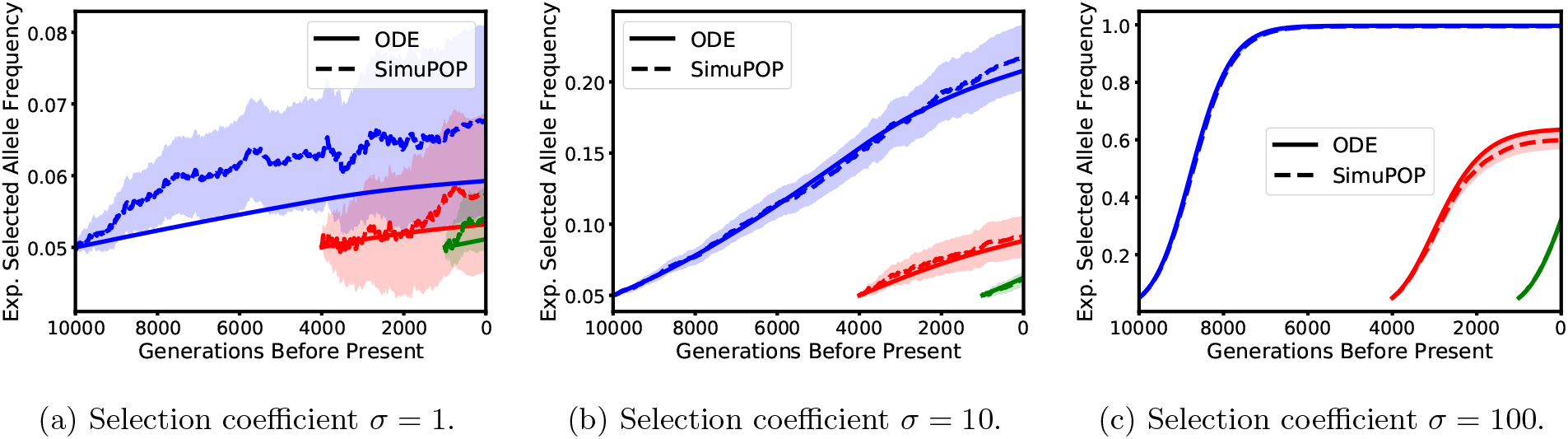
Expected trajectory of frequency of beneficial allele *A*. Blue, red, and green correspond to the dynamics starting 10,000 generations, 4,000 generations (beginning of the bottleneck), and 1,000 generations before present (beginning of exponential growth), respectively. The dotted and solid lines correspond to the simulation average and ODE solution, respectively, with the shaded region indicating a 95% confidence interval.

**Figure 5:**
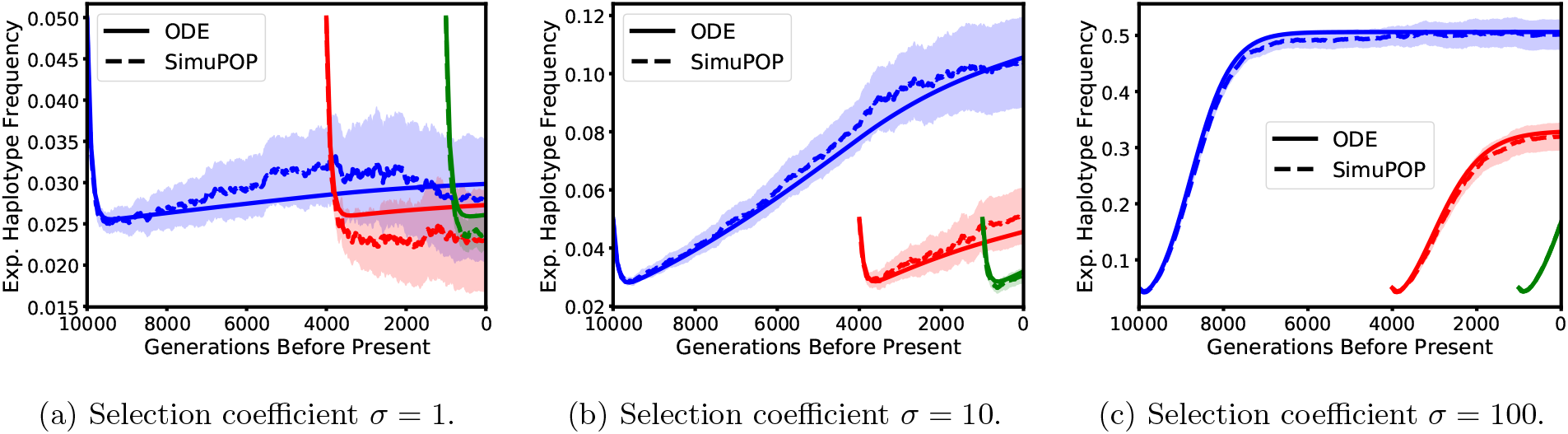
Expected frequency of the *AB* haplotype, carrying the beneficial allele. Again, blue, red, and green correspond to the dynamics starting 10,000 generations, 4,000 generations (beginning of the bottleneck), and 1,000 generations before present (beginning of exponential growth), respectively. The dotted and solid lines correspond to the simulation average and ODE solution, respectively, with the shaded region indicating the 95% confidence interval.

**Figure 6:**
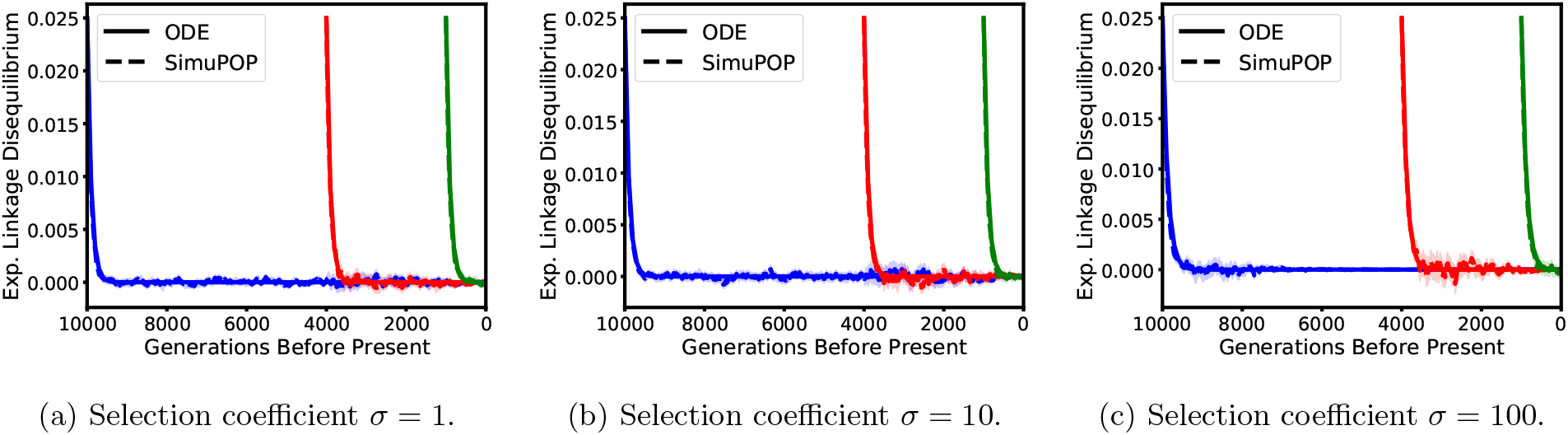
Expected linkage Disequilibrium between locus *A* and *B* for *ρ* = 400. Blue, red, and green correspond to the dynamics starting 10,000 generations, 4,000 generations (beginning of the bottleneck), and 1,000 generations before present (beginning of exponential growth), respectively. The dotted and solid lines correspond to the average of the simulations and ODE estimates, respectively, with the shaded region representing a 95% confidence interval.

**Figure 7:**
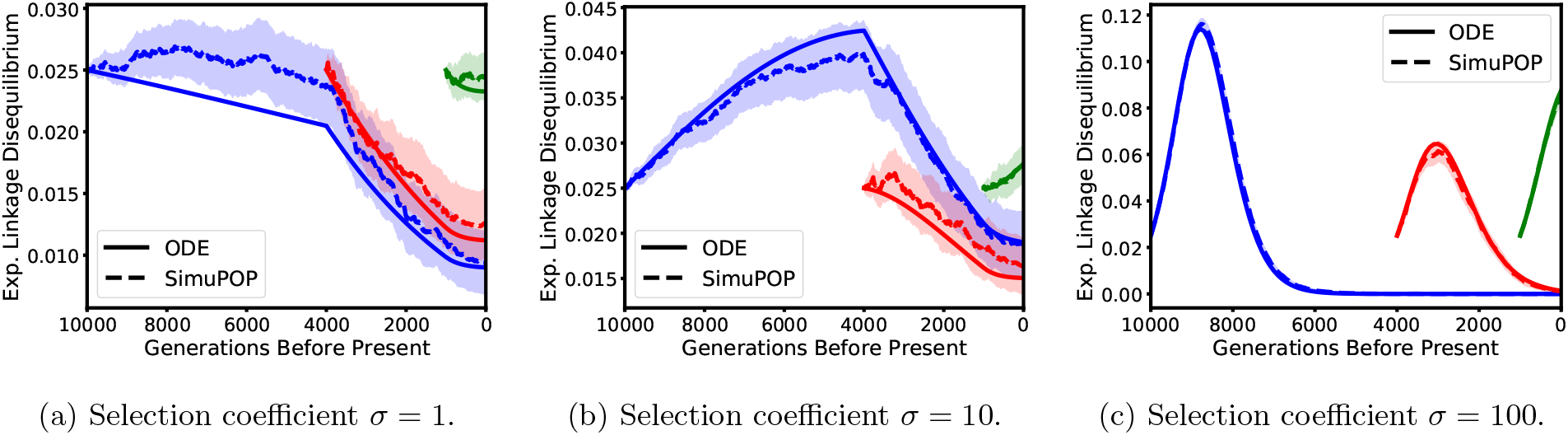
Linkage Disequilibrium between locus *A* and *B* for *ρ* = 0.0004. Blue, red, and green correspond to the dynamics starting 10,000 generations, 4,000 generations (beginning of the bottleneck), and 1,000 generations before present (beginning of exponential growth), respectively. The dotted and solid lines correspond to the average of the simulations and ODE estimates, respectively, with the shaded region representing a 95% confidence interval.

Next, in Figure 5, we show the trajectories of the expected frequency of haplotype AB, which carries the beneficial allele *A*. Here, we show the scenarios with a recombination rate of *ρ* = 400. This choice exhibits a broader spectrum of dynamics, since in the case *ρ* = 0.0004 linkage is very strong, and thus the dynamics are similar to the single locus case in Figure 4 (see Figure S1 in Supplementary Material). With *ρ* = 400 however, due to the strong recombination, we observe an initial decrease in the frequency of the *AB* haplotype, since the *A* allele is only introduced on the *B* background, and then quickly spreads to the *b* background as well. Subsequently, the dynamics are then again dominated by selection, and the haplotype *AB* increases, but only to a frequency of 0.5, since the *Ab* haplotype will also rise to that frequency in this scenario. Again, the increase is faster for stronger selection. To complement these dynamics, in Figure 6 we also depict the trajectory of the expected linkage disequilibrium (LD) between locus *A* and locus *B*:

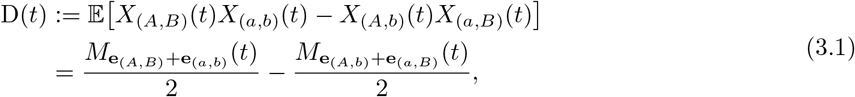

where **e**_*i*_ is the vector with 1 in component *i*, and 0 otherwise. We observe that it rapidly decreases, coinciding with the decrease of the expected frequency of the *AB* haplotype, but once it reaches zero, the haplotype frequencies starts increasing again.

Lastly, we also investigate the dynamics of the expected LD in the scenario with the lower recombination rate *ρ* = 0.0004, displayed in Figure 7. This lower recombination rate leads to much slower decay in LD, allowing us to exhibit the intricate interaction between selection, recombination, and demography. In the case of strong selection, we observe that LD can actually increase, the effect is however dependent on the population size. This behavior has already been noted before, see for example Kim and Nielsen (2004). As can be seen in Figure 7(a) and Figure 7(b), the rate of decrease changes substantially when the bottleneck at 4,000 generations before present and the exponential growth 1,000 generations before present is encountered. In addition to exhibiting the complex dynamics of the genetic variation in the population in these scenarios, the close match between the simulations and the numerical solutions demonstrates that our method accurately captures this dynamics and correctly accounts for the variable population size history.

In summary, we observe that our method captures the population dynamics of the expected frequencies and linkage disequilibrium well. Thus, the results present in this section provide evidence that our method efficiently and accurately computes the correct dynamics of the genetic variation at two-linked loci across a wide range of parameter settings, including selection at one of the loci.

#### 3.1.2 Initializing at Stationarity

We are furthermore interested in applying our method to study scenarios in which a novel beneficial allele arises in a population at stationarity, that is, the linked neutral allele is in mutation-drift equilibrium. Under the Wright-Fisher diffusion with recurrent mutation, this stationary distribution is given by a Beta(*θ, θ*) distribution (cf. Durrett, 2008, Sec. 7.5). In this case, the initial haplotype frequencies are not fixed values, but random and follow a certain distribution. Here, we initialize the ODEs with the moments of this distribution. Characterizing these moments requires additional assumptions about how the new mutation arises in the population. In particular, say the new beneficial allele arises at a low frequency *x_A_*. The new allele will fall either on the *B* or the *b* background. The probability of obtaining a sample containing both haplotype *AB* and *Ab* under this distribution is 0, and thus the only non-zero moments are those were either *n*_(*A,B*)_ = 0 or *n*_(*A,b*)_ = 0. In addition, the probability of *A* arising on the background *B* (or *b*) is proportional to the frequency of allele *B* in the population. As a result, the focal allele A tends to arise linked with the major allele more frequently than the minor allele. We give a detailed derivation of the initial moments for this scenario in Supplementary Material S2.4.2.

We chose the initial frequency for the beneficial allele as *x_A_* = 0.05 and initialize the ODEs using this distribution. This frequency is chosen to be 0.05 because, when initialized with fewer beneficial alleles, the selected allele is purged from the population most of the time, leading to very little difference from the neutral case for the expected quantities considered here. Computing the solution for different recombination rates allows us to explore how the impact of selection on linked neutral variation changes with increasing recombinational distance from the focal site under selection. Again, in the remainder of this section we compare the numerical solutions of the ODEs to the corresponding quantities obtained from simulations using `SimuPOP`. We assume the same demographic model as above and, assuming that two adjacent base-pairs are separated by a recombination distance of *ρ* = 0.0004, explore a 100 kbp region around the focal locus under selection. We consider 40 equally spaced neutral loci on either side of the selected side, the first of which is assumed to be perfectly linked (i.e. *ρ* = 0) with the selected locus and the outermost is separated by a recombination distance of *ρ* = 50,000 × 0.0004 = 20. In the `SimuPOP` simulations we simulate a population in which each individual has 41 loci, and for the numerical method, we solve the ODEs for each neutral locus paired with the selected locus, thus we solve it 40 times with different recombination rates. Note that we will mirror the plot in all cases so that the selected locus is in the center and it captures the dynamics of a 100 kbp region with a selected locus in the center. Since each of the 40 loci effectively represents 1,250 nucleotide sites, we also increase the population scaled mutation rate to *θ* = 0.625 in order to achieve the appropriate levels of neutral diversity. Finally, as in the previous section, we reduce the population size in the simulations by a factor of 10 to allow for tractable simulations and scale the parameters accordingly.

In Figure 8 we examine the expected heterozygosity at the neutral loci for different selection coefficients, defined as

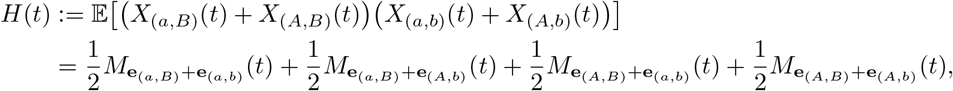

where *A* denotes the beneficial allele. The left-most column shows the results when the beneficial allele is introduced 10,000 generations before present (*η*_1_), thus experiencing the full demography, the middle column shows starting from just before the bottleneck (4,000 generations before present, *η*_2_), and the last column shows the results when starting at the beginning of the exponential growth (1,000 generations before present, *η*_3_). Note that in the examples where the beneficial is introduced at *η*_1_ and *η*_2_ the dynamics are initialized with the stationary distribution computed based on the larger (pre-bottleneck) population size. For the third column, in which the beneficial allele is introduced at *η*_3_ after 3,000 generations in a bottleneck, we use an approximation detailed in Supplementary Material S2.4.4 to compute the moments for the case when the beneficial allele is introduced into a population with a non-stationary distribution. Along the rows from top to bottom, the selection coefficients are *σ* = 1, 50, and 100. Note that in all cases, at the time the beneficial allele is introduced, the level of heterozygosity is the same across all neutral loci. As expected, this background level is lower when the beneficial allele is introduced after the bottleneck. Moreover, over time, selection reduces the heterozygosity at the selected locus, and this effect is stronger for larger selection coefficients. The neutral loci most closely linked to the selected locus also experience this reduction in heterozygosity, but the effect diminishes with increasing recombination distance, resulting in the characteristic “V” shape indicating a loss of genetic diversity near the selected locus, the hallmark signature of genetic hitchhiking. At the same time, the overall level of heterozygosity also decreases as the population enters the bottleneck. Moreover, in the case of strong selection, once the beneficial allele has swept to high frequency, the minimum of the “V” starts to increase again, and the signature of the selective sweep starts to erode. For completeness, we provide similar Figures for *x_A_* = 0.03 (Figure S4) and *x_A_* = 0.01 (Figure S5) in the Supplementary Material.

**Figure 8:**
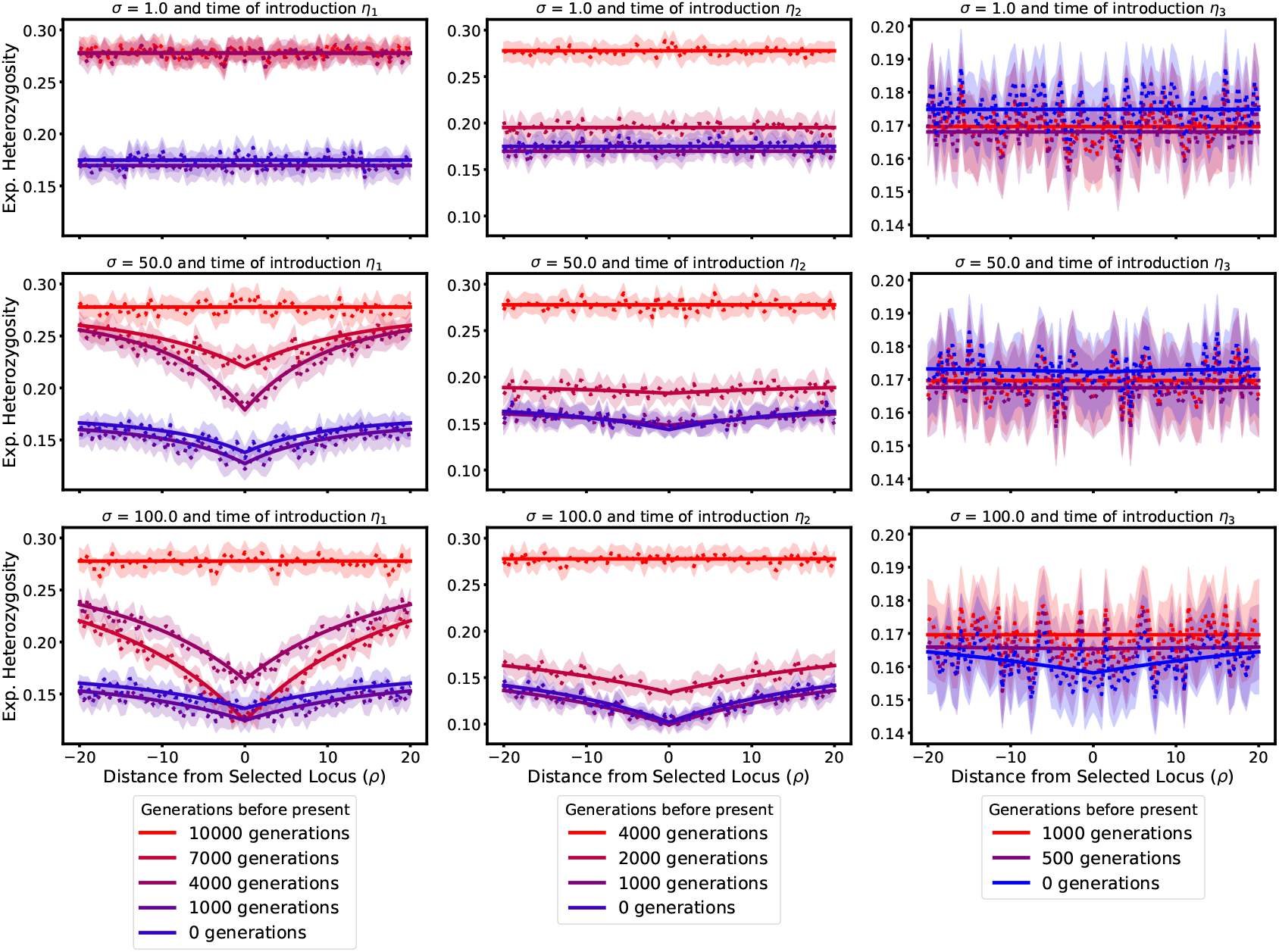
Expected heterozygosity across a 100 kbp region with different selection coefficients and times that the beneficial allele is introduced. The dotted and solid lines represent the simulations and numerical results, respectively, with the shaded region indicating a 95% confidence interval.

To complement these considerations on the dynamics of expected heterozygosity at the neutral sites, we also investigated a measure of LD in the same scenarios. Due to the stationarity assumption at the neutral site, regular LD at the time when the beneficial allele is introduced can be positive or negative with equal probability, and thus the expected LD as given in equation (3.1) is 0 at all later times. For this reason, we consider the expectation of LD squared, that is,

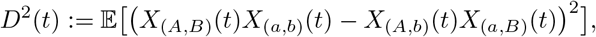

which can be expressed in terms of the moments of order 4. Figure 9 shows *D*^2^(*t*) in the same scenarios and for the same times as Figure 8. Since in the initial distribution, the beneficial allele arises on a given neutral background, that is, it is entirely linked to one neutral allele, at the time of introduction, we see a certain level of *D*^2^. At the loci that are far away from the focal locus, *D*^2^ goes to zero over time. However, closer to the selected locus, the dynamics of the selected allele causes *D*^2^ to increase strongly. For small selection coefficients, in contrast to little impact on the expected heterozygosity, we see a strong impact on *D*^2^. This is due to weak selection increasing the allele frequency continuously over long time intervals, and thus constantly building up LD. On the other hand, for strong selection, where the beneficial allele sweeps to fixation quickly, we observe a quick build-up of LD, but this signal also vanishes quickly once the beneficial allele is close to fixation. This build-up and decay has been exhibited in the literature before, for example by Zeng et al. (2021). For the strongest selection coefficient, we do not observe this build-up, since our resolution in time is likely not fine enough. For completeness, we provide similar Figures for *x_A_* = 0.03 (Figure S6) and *x_A_* = 0.01 (Figure S7) in the Supplementary Material.

**Figure 9:**
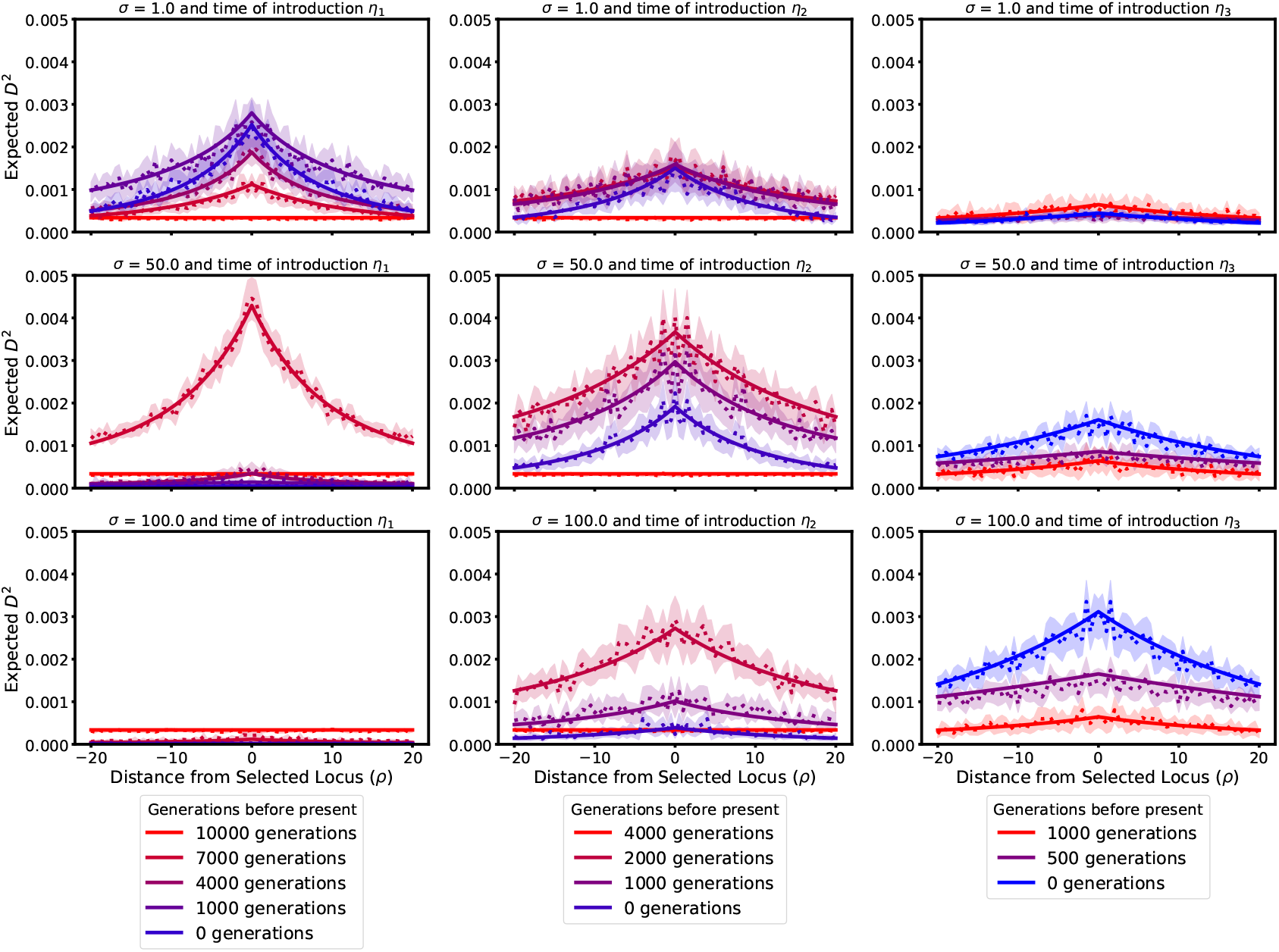
Expected *D*^2^ across a 100 kbp region with different selection coefficients and times that the beneficial allele is introduced. The dotted and solid lines represent the simulation and ODE results, respectively, with the shaded region representing a 95% confidence interval.

As Figure 8 and Figure 9 show, our method can be used to elucidate the extent of reduction in heterozygosity or inflation of linkage disequilibrium at certain recombination distances from the focal locus, and thus ultimately to characterize how wide the footprint of genetic hitchhiking extends in a variety of scenarios. Moreover, we observe again that the numerical solutions fall within the confidence bounds of the simulations, indicating that our numerical method is highly accurate in capturing the dynamics. In addition, the numerical solutions result in fairly smooth curves for the expected quantities of interest, whereas the curves estimated from the simulations exhibit a large degree of variability, which could only be reduced by performing many additional simulations, and would drastically increase computation time. In cases with much lower mutation rates than considered in our comparisons we would need many replicates of the simulations to even observe a single mutant at the neutral locus, resulting in even more variable estimates of very small quantities. However, the numerical solutions of the ODEs can readily incorporate the respective dynamics and yield accurate results at all scales.

### 3.2 Impact of Selection on Local Site-Frequency-Spectrum

In this section, we use our method to investigate the impact of selection at a focal locus on summary statistics of the neutral variation in a surrounding genomic window. Specifically, we will use the numerical solutions of the ODEs to efficiently and accurately compute the expectation of the site-frequency-spectrum (SFS) in this genomic window. We will consider only the full demographic model *η*_1_, see Figure 2, that is, the beneficial allele is introduced 10,000 generations before present. We use the non-recurrent mutation model and set the per locus mutation rate to *θ* = 0.0005 The recombination rate between adjacent sites is set to *ρ* = 0.0004. In order to compute the expected SFS for the whole window, we compute the two-locus moments using the ODEs over on a grid of 20 different recombination rates where the minimum is *ρ* = .0004, and the maximum is the maximum observed in the region of interest. As we will use window sizes 100 kbp, 500 kbp, and 1 Mbp, with the selected locus in the center, the maximum *ρ* is thus 20, 100, and 200, respectively, and the interior grid-points have logarithmic spacing. For a locus separated from the selected locus by a given recombination distance *ρ* that is not on the grid, we compute its vector of moments by interpolating between the moments computed on the grid. From these two-locus moments, which yield the two locus sampling probabilities, we can compute an individual locus’ contribution to the SFS by marginalizing over the selected locus which yields the vector of single-locus moments (sampling probabilities) at the neutral locus. Specifically, for a given sample size *n* and number of minor derived alleles *k*, we compute the sum of two-locus moments

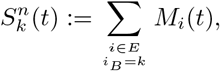

since *b* is the ancestral allele at the netrual site This yields the contribution to the *k*-th entry of the expected SFS by a neutral locus separated from the focal selected locus by the given recombination distance. Summing these quantities over all loci across the window of interest yields the full expected SFS.

We consider three different window sizes (100 kbp, 500 kbp, and 1 Mbp) in which the locus at the center is under genic selection with three different population scaled selection coefficients (*σ* = 1, 50, and 100), and compute both the expected SFS for a sample of size 31. Again, we note that the choice of this order strikes a balance between accuracy of the moment approximation technique and the size of the moment vector limiting computation. In principle, one could compute the expected SFS for larger samples, by computing the moments of a moderate order, and then using the solutions to approximate the higher orders using the moment approximation techniques outlined in Section 2.2. This does, however, require modifying the approximation technique at the boundaries, and this comes at a computational cost. We explore a preliminary implementation in Supplementary Material S4.

The initial frequency of the beneficial allele is *x_A_* = 0.05 and in each case and we use the non-recurrent mutation model to initialize the moments at the neutral loci, see Supplementary Material S2.4.3 for details. To validate the results we compare with simulations using `SLiM` (Haller and Messer, 2019; Haller et al., 2019), where the population size is again scaled down by a factor of 10 for efficiency. Each replicate undergoes 10,000 generations of burn-in so that the neutral loci are approximately at stationarity. The beneficial mutation is then introduced on a single chromosome chosen at random which is then copied onto 5% of the population. This allows us to initialize `SLiM` with the specified initial frequency for a beneficial allele linked to neutral background at stationarity. The simulation then undergoes the complete demographic history outlined in Figure 2, and we record the SFS at the indicated times, which are then averaged to get the expected value.

Figure 10 shows the expected SFS for the different window sizes and the different selection coefficients. At 10,000 generations before present, when the beneficial allele is introduced, the SFS exhibits the characteristic neutral shape, with an abundance of low frequency derived alleles and few high frequency derived alleles. Once the beneficial allele is introduced and subject to selection, we observe an increase in high frequency variants and a reduction of variants at intermediate frequency for strong selection. This is the expected pattern under genetic hitchhiking: To increase the frequencies at neutral loci linked to the beneficial allele. From the start of the bottleneck to the end, we observe a general reduction of genetic diversity, depleting alleles in all frequency classes, but affecting low frequency alleles more strongly. Once the exponential growth is encountered, we observe the characteristic inflation of low frequency minor alleles. The effects of selection are most pronounced for a small window size, and barely noticeable for large windows. This is to be expected, since the large windows contain a large number of neutral sites sufficiently removed from the selected locus to not be affected by the dynamics, and thus the neutral variation dominates the SFS. However, note that the total amount of variation in the window (see *y*-axis in the plots) is proportionately lower in the small windows. Thus, the SFS in the small window is more noisy when computed from real data, indicating that to characterize selection, one needs to strike a balance between large windows, with more statistical power in estimating the SFS, but less impact of selection, and small windows, where the impact of selection is greater, but fewer data points. Lastly, while the dynamics of the SFS in the given scenarios follows our expectation from population genetic principles, the fact that they are exhibited by our numerical solutions and match closely with the simulations shows again that our novel method indeed captures the relevant population genetic processes accurately. We also provide folded versions of these SFSs, computed using the recurrent mutation model, in Figure S8 in the Supplementary Material.

**Figure 10:**
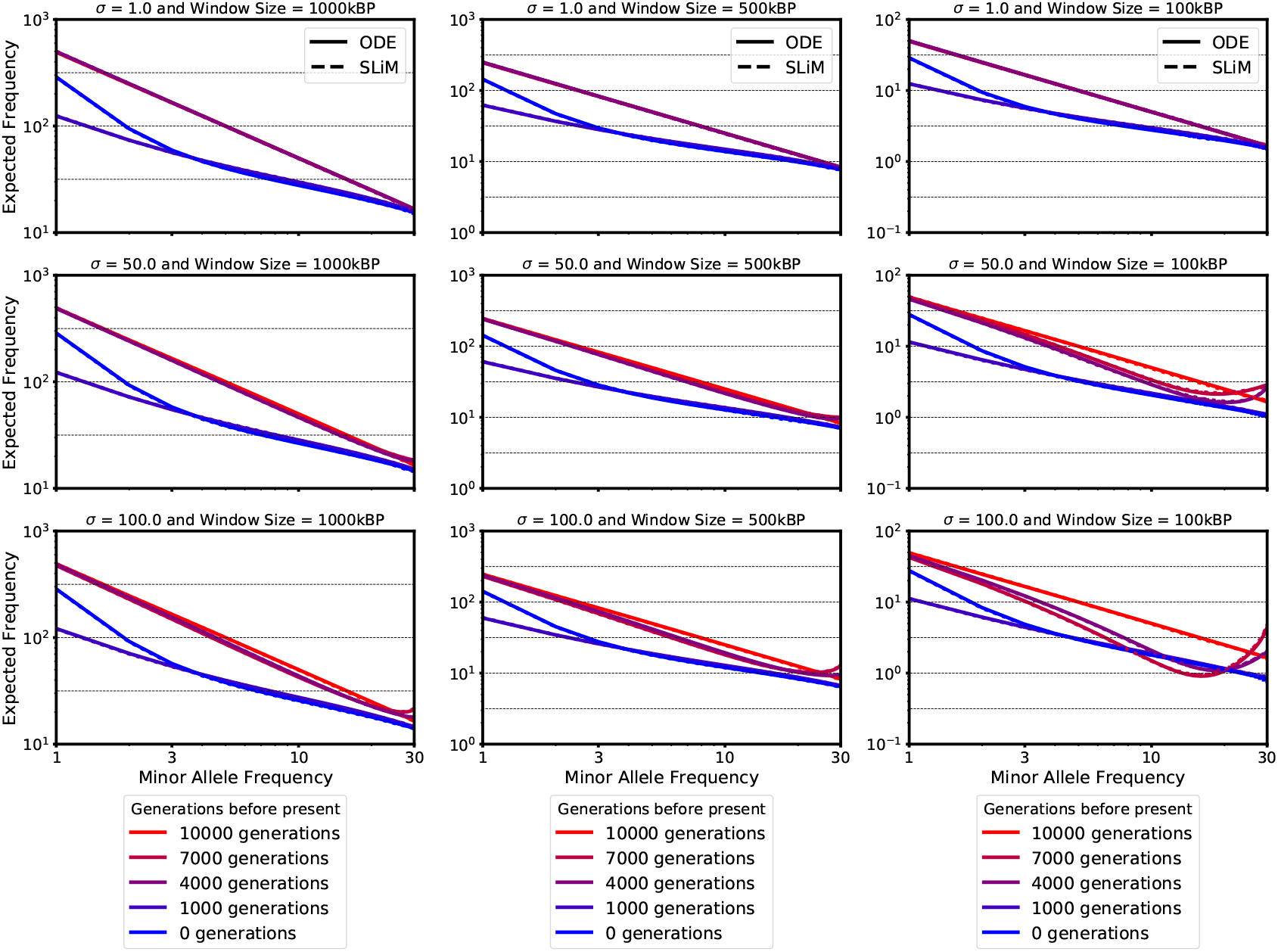
Local site-frequency-spectra for windows of size 100 kbp, 500 kbp, and 1 Mbp, for selection coefficients *σ* = 1, 50, and 100. The demographic history is given in Figure 2 and the beneficial allele is introduced 10,000 generations before present (*η*_1_). Both axis are on a log-scale. Note that for the third columns the y-axis limits have changed.

## 4 Discussion

We have presented a framework for computing moments of the transition density of the Wright-Fisher diffusion, or equivalently sampling probabilities, at two loci separated by an arbitrary recombination distance when selection is acting on the genetic variation. In addition, the framework accounts for arbitrary population size histories. We have explored our implementation of this framework in the case where genic selection is acting at one of the loci and demonstrated that it accurately captures the dynamics of relevant statistics under genetic hitchhiking in a demographic model that includes both a bottleneck and exponential growth, by comparing it to simulations using `SimuPOP` and `SLiM`. We believe this framework and our implementation enables progress in at least two directions.

On the one hand, it allows exploration of the impact of direct or linked selection on frequently used statistics computed from genomic data. As we have shown, the method can be used to compute a variety of quantities of interest under a range of demographic scenarios and varying strength of selection, more efficiently than using simulations. These quantities include expected allele or haplotype frequency trajectories, heterozygosity, and linkage disequilibrium. As patterns of LD are integral to a variety of existing population genetic tools, the framework described here provides a means of elucidating the impact of selection and demographic history on LD to allow for a better understanding of the patterns observed in genomic data. For example, in recent work by Ragsdale (2021), this framework has been applied to explore the impact of epistasis and dominance on shaping patters of signed LD observed in genic regions. As demonstrated in Section 3.1.1, our novel moment approximation technique is more accurate than the Jackknife approach employed by Ragsdale (2021). Thus, using this novel approximation technique in the analysis performed in Ragsdale (2021) and similar analyses has the potential to increase accuracy and substantiate the conclusions. In addition, the method can be used to compute the expected local SFS which allows for the exploration of linked and background selection on the SFS. Moreover, the outlined methodology provides a tractable means of calibrating the parameters for existing population genetics tools for detecting selection, for example, by elucidating the size of the genomic footprint under a given mode of selection and population size history.

On the other hand, our numerical framework is a key building block for developing likelihood-based tools to estimate the location, strength, and mode of selection from population genomic data, while accounting for arbitrary demographic scenarios. Including neutral variation around the selected locus into the likelihood model, has increased power over single-locus methods (Pavlidis and Alachiotis, 2017; Hejase et al., 2020). A possible approach is to use the two locus sampling probabilities computed using our method in a composite likelihood framework similar to the one introduced by McVean et al. (2002), that is, multiplying the likelihoods for each locus in a window paired with the candidate selected locus. Such a framework would allow one to compute maximum likelihood estimates of selection coefficients in general models of selection, and apply likelihood-based tests to determine significance.

We want to emphasize that the current implementation of our numerical method, while having many similarities with the method introduced by Ragsdale (2021), aims at a different area of application. The implementation by Ragsdale (2021) is designed to study aggregate patterns of two-locus LD genome-wide or in specific classes of genomic variation. The method implements a two-locus extension of the infinite sites model, and thus the entries of the moment vector are the expected value of the number of times a certain haplotype configuration is observed. Furthermore, Ragsdale (2021) exclusively use the two-locus stationary distribution as initial condition. Our numerical framework presented here, on the other hand, aims at describing the probabilities of observing a certain configurations in a sample of haplotypes at two specific loci. It can thus also be used to describe neutral genetic variation around a specific locus under selection in a certain genomic window. To this end, we implemented the recurrent mutation model, and a two-locus version of a non-recurrent mutation model. We also derived an initial condition where a novel beneficial allele is linked to neutral background variation. While both methods can, in principle, be modified to either of these scenarios, the choice of a user as to which implementation should be preferred is currently primarily based on the application domain they are interested in.

Our framework and its implementation can be extended into several useful directions. Here, we focused on describing how neutral genetic variation in a genomic region is affected by a single selective sweep. Another promising application of the framework is to model background selection, where neutral variation is effected by recurrent selective sweeps on deleterious variation that arises in the genomic background. Such scenarios have been studied in the literature, under constant population size (Charlesworth et al., 1993, 1995; Hudson and Kaplan, 1995; Nordborg et al., 1996; Nicolaisen and Desai, 2013), varying population size (Zeng, 2013), and in subdivided populations (Zeng and Corcoran, 2015; Ewing and Jensen, 2016). Our framework will be useful to extend and complement these results.

Moreover, implementing functionality for general modes of diploid selection requires no additional theoretical developments. For example, models of balancing selection have received much attention in the literature (e.g. Zhao and Charlesworth, 2016; Zeng, 2013). However, since general diploid selection requires increasing the order of the approximated moments by two, additional testing needs to be performed in order to ensure its accuracy. Nonetheless, we do expect our novel approximation scheme to generalize well to this case. Moreover, we have presented our framework in the setting of a single panmictic population, accounting for arbitrary population size history. It is natural to consider extending the framework to multiple populations with an arbitrary demographic history. This would enable applying it in situations were a beneficial alleles arises in only one population, and contrasting the genetic variation between populations, to study ancestral selection using the signatures from multiple extant population, or to study selection on introgressed genetic variation. Extending the theoretical framework to multiple populations is possible, since Ragsdale and Gravel (2019) do use multi-population moments of lower order to estimate demographic parameters under neutrality. However, extensions of the framework including selection present additional challenges for the implementation. Obtaining an accurate approximation of the dynamics under selection necessitates solving the ODEs for moments of sufficient high order to ensure the accuracy of the moment approximation when “closing” the ODEs. A straightforward extension would necessitate computing moments of high order in all populations, and thus the dimensionality equals the number of single population moments exponentiated by the number of populations. Since this number quickly grows to become infeasible, additional approximations will be required. A potential approach could be to compute the single locus dynamic for the selected allele first using higher order moments, and then “attach” the dynamics of lower order moments for the linked neutral loci in a second step. Other approaches could include reducing the ODEs from a system that keeps track of all moments, to a system where only bulk quantities, like sums of moments, are considered. Ultimately, the framework and implementation introduced in this paper present a promising foundation upon which to build these applications and extensions.

## Software Availability

A python implementation of the numerical procedure as well as scripts to create the figures in this manuscript can be found at https://github.com/steinrue/two_locus_selection_moments.

## Acknowledgements

We thank Aaron Ragsdale, Simon Gravel, and the population genetics community in Chicago for many inputs and helpful feedback on the methodology and manuscript. Moreover, we thank two anonymous reviewers, whose comments and suggestions have substantially enriched the manuscript.

## A Appendix: Derivation of Moments

### A.1 Drift

Recall the term of the generator corresponding to genetic drift, 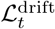, defined in (2.2). Then the contribution of genetic drift to the time derivative of moment *M*_n_(*t*) is,

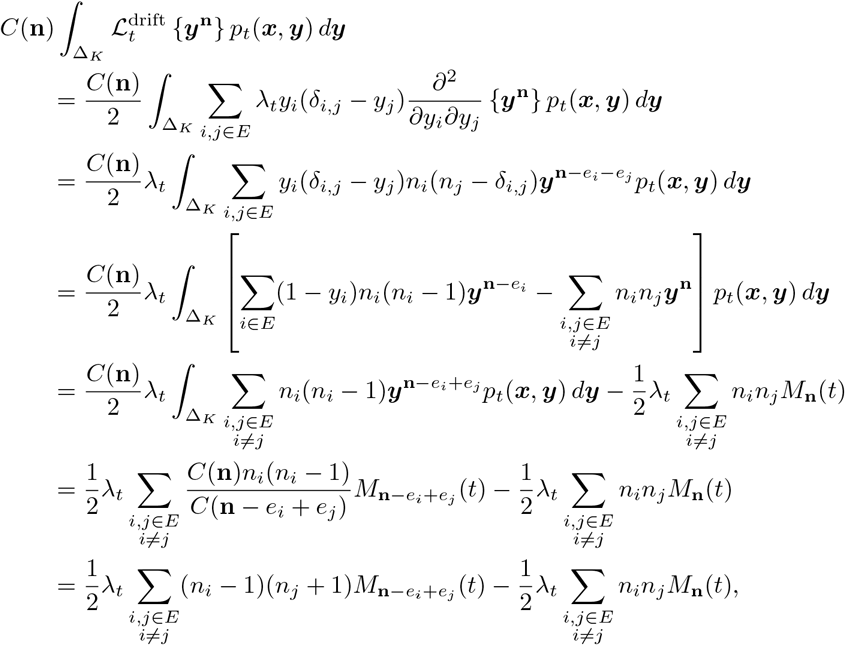

where *e_i_* denotes a vector with all components 0, except the *i*-th entry, which is equal to 1. It follows that we can define 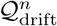 as follows,

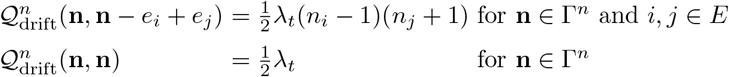

with all other entries being zero.

### A.2 Mutation

#### A.2.1 Recurrent Mutation

Recall the term of the generator corresponding to mutation, 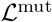, defined in (2.3). Then the contribution of mutation to the time derivative of moment *M*_n_(*t*) is,

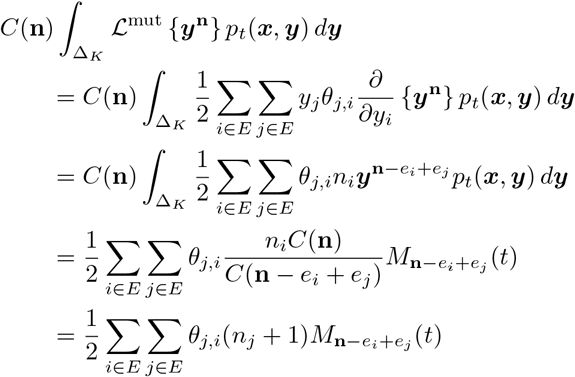

From this, we can define 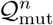 as follows,

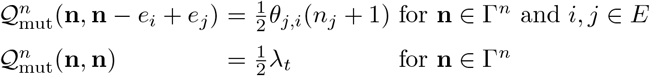

with all other entries being zero.

#### A.2.2 Non-recurrent Mutation

To model non-recurrent mutation, we closely follow the derivation of Ragsdale and Gravel (2019), specifically Equations (S19) to (S24) in their supplement. For simplicity, we assume a two-locus two-allele model where the first locus, with allele *a* and *A*, is under selection and the second locus, with alleles *b* and *B*, is neutral. We designate *a* and *b* as the ancestral alleles. We furthermore assume that mutations only occur at the neutral locus at rate 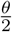. Moreover, in a non-recurrent mutation model, mutations can only change the ancestral allele into the derived allele, and thus, mutations only occur in haplotype configurations where all alleles are ancestral, as shown, for example, by Evans et al. (2007), Jouganous et al. (2017), or Ragsdale and Gravel (2019).

However, our implementation has a key difference: Ragsdale and Gravel (2019) use an infinite sites model, where each entry of the moment vector corresponds to an expected number of pairs of sites with a given configuration. Consequently, they inject mutations into configurations with one derived allele, and do not explicitly keep track of all configurations with only ancestral alleles. Since we are considering moments that represent sampling probabilities, they do have to sum to 1. If we inject mass into configurations with one derived allele, then we have to remove this mass from the configurations with all ancestral alleles, which we explicitly track. We then follow Ragsdale and Gravel (2019) and similar approaches using the infinite-sites model (Evans et al., 2007; Jouganous et al., 2017), and inject mass through purely inhomogeneous terms at a total rate of 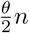 at the neutral locus. This total rate is then distributed proportionally to the different configurations with one derived allele, similar to Equations (S19) to (S24) in the supplement of Ragsdale and Gravel (2019).

To specify the rates for the non-recurrent mutation model, we first compute the total probability mass of configurations with all ancestral alleles at the neutral locus as

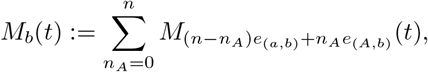

and the relative mutation rate for each such configuration as

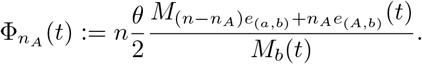

Then, the contribution of non-recurrent mutation to the derivative of the moment vector with respect to *t* is

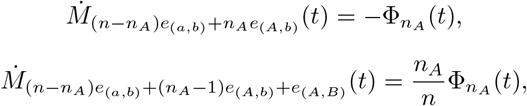

and

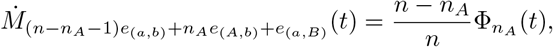

for *n_A_* ∈ {0,…, *n*}. We use this implementation to obtain the results depicted in Figure 10.

### A.3 Selection

Recall the term of the generator corresponding to mutation, 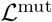, defined in (2.4). Then the contribution of natural selection to the time derivative of moment *M*_n_(*t*) is,

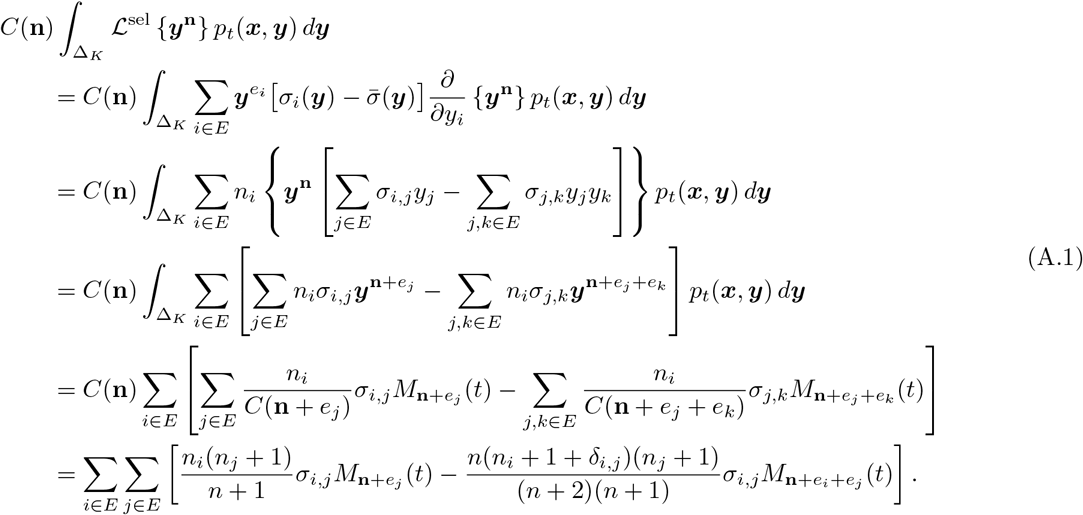

It then follows that 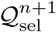 is defined as follows,

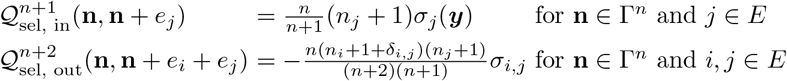

with all other entries being zero. Since moments of order *n* +1 can be computed from the moments of order 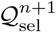 can be re-written so that it maps moments of order *n* + 2 to those of *n*. Namely, denoting the new matrix as 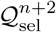, it can be expressed as 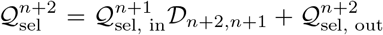, where 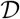 is the downsampling matrix, see Supplementary Material S2.2 for more details.

#### A.3.1 Genic Selection

Consider the case of genic selection at locus *A*. Namely, for all *i, j* ∈ *E*, define *σ_i,j_* as,

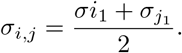

From the fourth line of (A.1) we have,

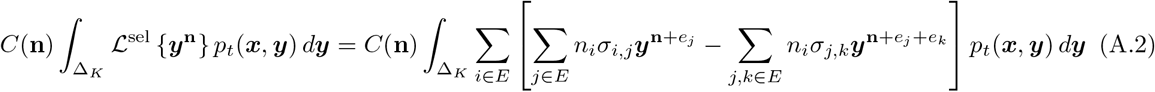

The first term on the RHS of the above equation can be written as,

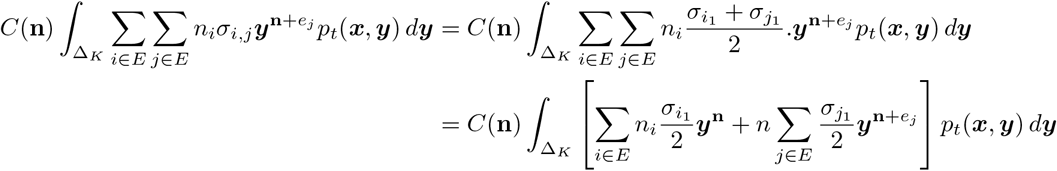

Then the second term on the RHS of (A.2) can be written as,

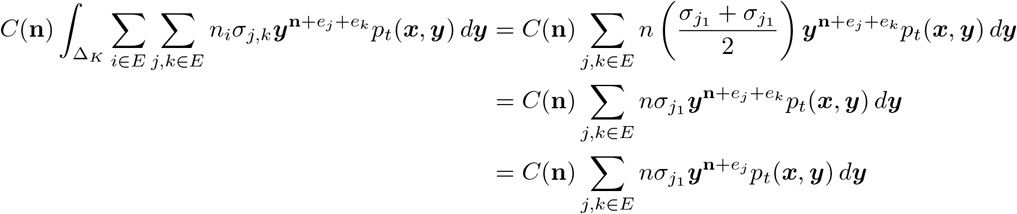

where the first equality follows from the definition of *σ_j,k_*, the second follows from the symmetry of *j* and *k*, and the third follows from the fact that 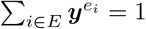. Substituting this back into equation (A.2) yields,

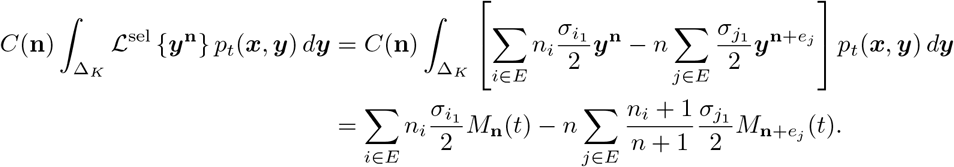

It then follows that 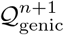 is defined as follows,

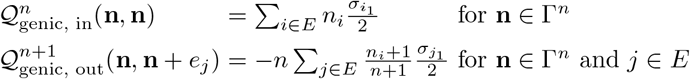

with all other entries being zero. As with the case of general selection 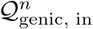 can be written as 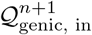 in and combined with 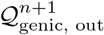 to get 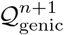.

### A.4 Recombination

Recall the term of the generator corresponding to recombination, 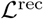, defined in (2.5). Then the contribution of recombination to the time derivative of moment *M*_n_(*t*) is,

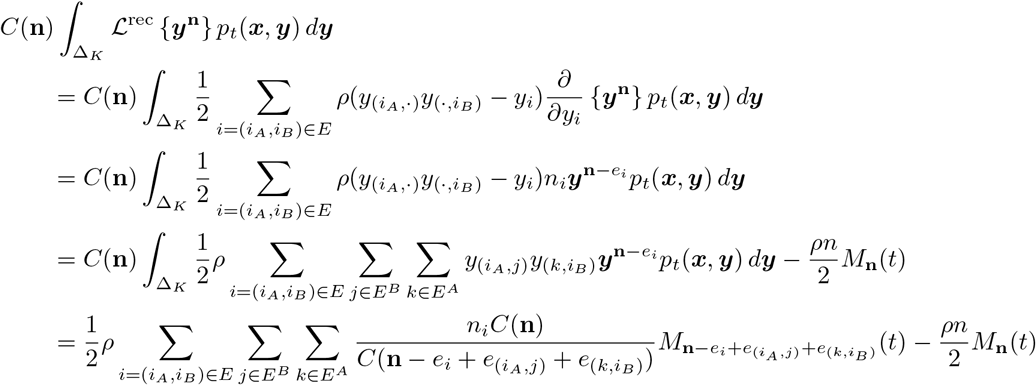

The quantity 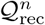 can then be defined as follows,

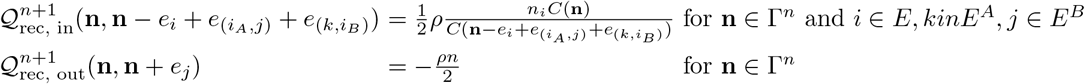

with all other entries being zero. As in the case of selection 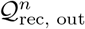 can be written as 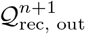 and combined with 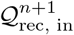 to get 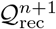.

