## Supplementary Material for "A numerical framework for genetic hitchhiking in populations of variable size"

#### S1 Proof of Lemma 2.3 in the Main Text

*Proof of Lemma 2.3.* Recall the probability measure

$$\mu := f(y)dy$$

on $[0,1]$ with density $f(\cdot)$ with respect to the Lebesgue measure $dy$, and the sequence of probability measures

$$\mu^n := \bar{f}_n(y)dy$$

with piece-wise constant densities

$$\bar{f}_n(y) := \sum_{i=0}^{n} \Big( \int_0^1 b_i^n(z) f(z) dz \Big) (n+1) \mathbb{1}_{[\frac{i-0.5}{n}, \frac{i+0.5}{n})}(y)$$

with $y \in [0,1]$, where

$$b_i^n(z) = \binom{n}{i} z^i (1-z)^{n-i}$$

are the Bernstein polynomials. To show weak convergence of $\mu_n$ to $\mu$, consider a bounded, Lipschitz continuous function $g : [0,1] \to \mathbb{R}$, that is, there exists a $K > 0$ such that

$$\big|g(x) - g(y)\big| \leq K|x-y|$$

holds for all $x, y \in [0,1]$. Moreover, let $g_{\max} \geq |g(\cdot)|$ be a bound on the function. Then, we have to show that

$$\int_0^1 g\mu^n \to \int_0^1 g\mu$$

for all such $g(\cdot)$. To this end, note that

$$\int_0^1 g\mu = \int_0^1 g(y) f(y) dy$$

holds. Furthermore, for the approximating sequence $\mu_n$, we have

$$
\begin{aligned}
\int_0^1 g\mu^n &= \int_0^1 g(y)\bar{f}_n(y)dy \\
&= \sum_{i=0}^{n}\Big(\int_0^1 b_i^n(z)f(z)dz\Big)(n+1)\int_0^1 \mathbb{1}_{[i-\frac{1}{2n},i+\frac{1}{2n})}(y)g(y)dy \\
&\le \sum_{i=0}^{n}\Big(\int_0^1 b_i^n(z)f(z)dz\Big)(n+1)\Big(\int_{i-\frac{1}{2n}}^{\frac{i}{n}}\Big(g\Big(\frac{i}{n}\Big)+K\Big(\frac{i}{n}-y\Big)\Big)dy+\int_{\frac{i}{n}}^{i+\frac{1}{2n}}\Big[g\Big(\frac{i}{n}\Big)+K\Big(y-\frac{i}{n}\Big)\Big]dy\Big) \\
&= \sum_{i=0}^{n}\Big(\int_0^1 b_i^n(z)f(z)dz\Big)2(n+1)\int_0^{\frac{1}{2n}}\Big(g\Big(\frac{i}{n}\Big)+Ky\Big)dy \\
&= \sum_{i=0}^{n}\Big(\int_0^1 b_i^n(z)f(z)dz\Big)\frac{n+1}{n}\Big(g\Big(\frac{i}{n}\Big)+K\frac{1}{4n^2}\Big) \\
&= \sum_{i=0}^{n}\Big(\int_0^1 b_i^n(z)f(z)dz\Big)g\Big(\frac{i}{n}\Big)\frac{n+1}{n}+K\frac{n+1}{4n^2},
\end{aligned}
$$

where the inequality follows from the Lipschitz continuity of $g(\cdot)$ and setting $x=\frac{i}{n}$, and the last equality holds, because

$$\sum_{i=0}^{n}\Big(\int_0^1 b_i^n(z)f(z)dz\Big)=1.$$

We can proceed along the same lines to obtain a lower bound which ultimately yields

$$
\begin{aligned}
\int_0^1 g\mu^n &= \frac{n+1}{n}\sum_{i=0}^{n}\Big(\int_0^1 b_i^n(z)f(z)dz\Big)g\Big(\frac{i}{n}\Big)+O\big(n^{-1}\big) \\
&= \frac{n+1}{n}\int_0^1\sum_{i=0}^{n} b_i^n(z)g\Big(\frac{i}{n}\Big)f(z)dz+O\big(n^{-1}\big)
\end{aligned}
\tag{S1.1}
$$

The Weierstrass approximation theorem (e.g. Klenke, 2008, Ex. 5.15) applied to the Bernstein polynomials $b_i^n(z)$ can be used to show that

$$\sum_{i=0}^{n} b_i^n(z)g\Big(\frac{i}{n}\Big)\to g(z)$$

uniformly on $[0,1]$ as $n\to\infty$. Furthermore, this expression can be bounded by

$$\sum_{i=0}^{n} b_i^n(z)g\Big(\frac{i}{n}\Big)\le g_{\max}\sum_{i=0}^{n} b_i^n(z)=g_{\max}$$

for all $x$, and thus

$$\int_0^1 g\mu^n\to\int_0^1 g(y)f(y)dy \tag{S1.2}$$

follows from the dominated convergence theorem (Klenke, 2008, Cor. 6.26) applied to the integral on the right hand side of equation (S1.1). Convergence of the integrals in (S1.2) for all bounded Lipschitz continuous functions $g(\cdot)$ is equivalent to the weak convergence of $\mu_n$ to $\mu$ due to the Portemanteau theorem (Klenke, 2008, Thm. 13.16).

□

### S2 Implementation

Our method relies on numerically solving the ODE (2.7) in the main text. This is ultimate accomplished using a modified Runge-Kutta-Fehlberg 4(5) method. The two main numerical challenges faced were to (1) quickly and accurately estimate moments of order $n+1$ and $n+2$ from those of order $n$, and then ensuring that the numerical solution to (2.7) remains stable while using the approximation. In this section we will detail how the method is numerically implemented.

#### *S2.1 ODE Solver Implementation*

We solve the ODE numerically using a modified Runge-Kutta-Fehlberg 4(5) method (e.g. Kincaid et al., 2009, Ch. 8.3). Since the state-space is a simplex, the implementation needs to take these constraints into account. In particular, at each step, all negative values are set to zero and values greater than one are set to one. Then, the state-vector is renormalized so that all the components sum to one.

#### *S2.2 Parimonious Moments*

We now introduce a definition which is useful for the implementation of the ODE. First, consider the following identity:

$$
\begin{aligned}
\mathbf{M_n} &= C(\mathbf{n}) \int_{\Delta_K} \boldsymbol{y}^{\mathbf{n}} dp(\boldsymbol{y}) = C(\mathbf{n}) \int_{\Delta_K} \boldsymbol{y}^{\mathbf{n}} \left( \sum_{i=1}^{K} y_i \right) dp(\boldsymbol{y}) = C(\mathbf{n}) \int_{\Delta_K} \left( \sum_{i=1}^{K} \boldsymbol{y}^{\mathbf{n}+e_i} \right) dp(\boldsymbol{y}) \\
&= C(\mathbf{n}) \int_{\Delta_K} \left( \sum_{i=1}^{K} \boldsymbol{y}^{\mathbf{n}+e_i} \right) dp(\boldsymbol{y}) = \sum_{i=1}^{K} \frac{C(\mathbf{n})}{C(\mathbf{n}+e_i)} \mathbf{M}_{\mathbf{n}+e_i}.
\end{aligned} \tag{S2.1}
$$

This shows that every moments of order $n$ can be explicity computed as a linear combination of moments of order $n+1$. One can apply this identity multiple time to represent $\mathbf{M}_n$ in terms of $\mathbf{M}_{n+m}$ for any $m > 0$. If the two vectors $M_n$ and $M_{n+m}$ satisfy these identities for all components of $M_n$ we say that they are "parsimonious". Furthermore, one can use (S2.1) to define a sparse matrix $\mathcal{D}_{n+m,n}$ which maps $\mathbf{M}_{n+m}$ to $\mathbf{M}_n$.

#### *S2.3 Evaluation of Derivative*

The time-derivative of the state variable $\mathbf{M}_{n+1}(t)$ is composed of four components (genetic drift, mutation, recombination, selection). For convenience, we split recombination and selection into *in* (positive term) and *out* (negative term). Each of the six terms can then be computed by multiplying the moment vector with a sparse matrix:

1. Net flux due to genetic drift.
2. Net flux due to mutation.
3. Flux in due to selection.
4. Flux out due to recombination.
5. Flux out due to selection.
6. Flux in due to recombination.

Terms 1-4 can be expressed as linear combinations of the order $n$ moments while 5 and 6 need moments of order $n+1$ (or $n+2$ in the case of general diploid selection) and rely on the moment approximation. However, we noticed that in most practical scenarios the ODE is more accurate and stable when using the moments of order $n$ which are parsimonious with those moments of order $n+1$. When evaluating the derivative, we thus employ the vector of moments which are parsimonious with the estimated $M_{n+1}$ rather than the original $M_n$ for 3 and 4. Furthermore since $\mathcal{D}_{n+1,n}$ is sparse this can be implemented efficiently.

#### *S2.4 Initial Distribution*

In this section we outline how the initial moments for the ODEs are computed. We consider two types of initial conditions. In the first, the population starts from a fixed haplotypes frequency $x = [x_{AB}, x_{Ab}, x_{aB}, x_{ab}]$. This case is straightforward as the moments can easily be obtained using a multinomial distribution. In the second case, we assume that the neutral locus $B$ is initialized from a given distribution. The beneficial allele at locus $A$ is then introduced at a given frequency on this background. In the main text we consider both stationary and non-stationary distributions for the neutral locus.

##### S2.4.1 Given Distribution at Neutral locus

Suppose we model the case in which a selected allele is introduced at frequency $x_A$ at time 0 while the neutral site is has a distribution with density $p_B$. A natural way to implement this seems to just initialize the Wright-Fisher diffusion with the probability measure $\delta_{x_A} \times p_B$. However, this measure is not correct, especially if one is interested in modeling the impact of selection on nearby neutral sites. The problem is that it is modeling a situation in which the new allele $A$ can land on both the $B$ and $b$ background simultaneously. To see this note that if we were to take $\delta_{x_A} \times p_B(x)dx$ with $x_A = 1/2N$ (corresponding to a single selected allele) and computed the sampling probabilities for a sample of size 2, the probability of sampling one $AB$ haplotype and one $Ab$ haplotype would be non-zero. Instead the initial distribution should be

$$\begin{aligned} &I(1 - x_{AB} \geq x_B \geq x_{AB})(x_B \delta_{(x_A,0)}(x_{AB}, x_{Ab}) + (1 - x_B)\delta_{(0,x_A)}(x_{AB}, x_{Ab})) \times p_B(x_B)dx_B \\ &\quad + I(1 - x_A < x_B)\delta_{(x_A,0)}(x_{AB}, x_{Ab}) \times p_B(x_B)dx_B \\ &\quad + I(x_B < x_A)\delta_{(0,x_A)}(x_{AB}, x_{Ab}) \times p_B(x_B)dx_B, \end{aligned} \tag{S2.2}$$

where $I(\cdot)$ is the indicator function that is 1 if the condition is true, and zero otherwise. Under this joint distribution, the marginal distribution of alleles at the neutral locus follows a distribution with density $p_B$. Furthermore, the distribution of the frequency of the allele $A$ is a point mass at frequency $x_A$. This reflects that if one were to randomly introduce a mutation $A$, it has probability $x$ of landing on the same chromosome as a $B$ and $1 - x$ of arising on the $b$ background. The indicator functions reflect the fact that the frequencies of $B$ and $b$ cannot be smaller than the frequencies of $AB$ and $Ab$, respectively. In particular, the first line represents the case in which the frequency of $B$ and $b$ is above that of $A$ and thus both $AB$ and $Ab$ are possible. The second (resp. third) line correspond to the case in which $b$ (resp. $B$) is too small for $Ab$ (resp. $Ab$) to exist. Note that this measure is discontinuous as $x_B$ increases across the threshold $x_A$. This is due to the fact that the marginal distribution $p_B(x_B)$ is assuming a population of infinite size, whereas introducing an allele in a finite number of individuals is assuming a finite population. Indeed the probability measure (S2.2) is an approximation of both, however it approximates the scenario sufficiently well for small $x_A$. Given a moment configuration, the moment can be computed by integrating the appropriate multinomial probability-mass-function against the measure (S2.2). This can be split of up into three integrals which correspond to the cases $x < x_A$, $x_a \leq x \leq 1 - x_A$, and $x < 1 - x_A$, respectively:

$$\begin{aligned} M_n(t) = \binom{n}{n_{AB}, n_{Ab}, n_{aB}, b_{ab}} \Bigg[ &\int_0^{x_A} 0^{n_{AB}} (x_A)^{n_{Ab}} x^{n_{aB}} (1 - x - x_A)^{n_{ab}} p_B(x)dx \\ &+ \int_{x_A}^{1-x_A} \Big( (x_A)^{n_{AB}} 0^{n_{Ab}} (x - x_A)^{n_{aB}} (1 - x)^{n_{ab}} x \\ &\qquad + 0^{n_{AB}} (x_A)^{n_{Ab}} x^{n_{aB}} (1 - x - x_A)^{n_{ab}} (1 - x) \Big) p_B(x)dx \\ &+ \int_{1-x_A}^{1} (x_A)^{n_{AB}} 0^{n_{Ab}} (x - x_A)^{n_{aB}} (1 - x)^{n_{ab}} p_B(x)dx \Bigg]. \end{aligned} \tag{S2.3}$$

Now, we consider the case of the stationary (Beta) distribution under recurrent mutation, the stationary distribution under the non-recurrent mutation model, and a set of moment taken from a non-stationary distribution separately:

##### S2.4.2 Stationary Beta Distribution

We first consider the case of the neutral allele starting from the stationary distribution of the Wright-Fisher diffusion with recurrent mutation. Namely, suppose $p_B$ is the density of a Beta$(\beta_1, \beta_2)$ distribution where $\beta_1$ and $\beta_2$ are the population scaled mutation rates at locus $B$. Considering the last integral in equation (S2.3) and assuming $n_{Ab} = 0$

$$\begin{aligned}
&\int_{1-x_A}^{1} (x_A)^{n_{AB}} (x - x_A)^{n_{aB}} (1-x)^{n_{ab}} p_B(x) dx \\
&= \int_{1-x_A}^{1} (x_A)^{n_{AB}} (x - x_A)^{n_{aB}} (1-x)^{n_{ab}} \frac{x^{\beta_1 - 1}(1-x)^{\beta_2 - 1}}{B(\beta_1, \beta_2)} dx \\
&= \int_{1-x_A}^{1} (x_A)^{n_{AB}} \sum_{i=0}^{n_{aB}} \left[ \binom{n_{aB}}{i} (-x_A)^{n_{aB}-i} \frac{x^{\beta_1+i-1}(1-x)^{\beta_2+n_{ab}-1}}{B(\beta_1, \beta_2)} \right] dx \\
&= \sum_{i=0}^{n_{aB}} \binom{n_{aB}}{i} (-x_A)^{n_{aB}-i} (x_A)^{n_{AB}} \frac{B(\beta_1 + i, \beta_2 + n_{ab}) - B(1 - x_A; \beta_1 + i, \beta_2 + n_{ab})}{B(\beta_1, \beta_2)}
\end{aligned} \tag{S2.4}$$

where $B(\cdot, \cdot)$ and $B(\cdot; \cdot, \cdot)$ are the Beta function and incomplete Beta function, respectively. By a similar argument the first term equals,

$$\begin{aligned}
&\int_{0}^{x_A} (x_A)^{n_{Ab}} x^{n_{aB}} (1 - x - x_A)^{n_{ab}} (1-x) p_B(x) dx \\
&= \sum_{i=0}^{n_{ab}} \binom{n_{ab}}{i} (-x_A)^{n_{ab}-i} (x_A)^{n_{Ab}} \frac{B(x_A; \beta_1 + n_{aB}, \beta_2 + i)}{B(\beta_1, \beta_2)}
\end{aligned} \tag{S2.5}$$

for $n_{AB} = 0$. Finally the middle term equals:

$$\begin{aligned}
&\int_{x_A}^{1-x_A} \Big[ (x_A)^{n_{AB}} 0^{n_{Ab}} (x - x_A)^{n_{aB}} (1-x)^{n_{ab}} x + 0^{n_{ab}} (x_A)^{n_{Ab}} x^{n_{aB}} (1 - x - x_A)^{n_{ab}} \Big] p_B(x) dx \\
&= 0^{n_{Ab}} \sum_{i=1}^{n_{aB}} \binom{n_{aB}}{i} (-x_A)^{n_{aB}-i} (x_A)^{n_{AB}} \frac{B(1 - x_A; \beta_1 + i + 1, \beta_2 + n_{ab}) - B(x_A; \beta_1 + i + 1, \beta_2 + n_{ab})}{B(\beta_1, \beta_2)} \\
&+ 0^{n_{AB}} \sum_{i=1}^{n_{ab}} \binom{n_{ab}}{i} (-x_A)^{n_{ab}-i} (x_A)^{n_{Ab}} \frac{B(1 - x_A; \beta_1 + n_{aB}, \beta_2 + i + 1) - B(x_A; \beta_1 + n_{aB}, \beta_2 + i + 1)}{B(\beta_1, \beta_2)}.
\end{aligned} \tag{S2.6}$$

##### S2.4.3 Non-recurrent Mutation

In the case where the neutral locus is initialized using the stationary measure of the non-recurrent mutation model, we use $p_B(x) = \frac{\theta}{x}$, where $\theta$ is the population-scaled mutation rate. Note that this is not a probability measure since it does not integrate to 1, but we can substitute it into equation (S2.3) regardless. We obtain the same expressions in equations (S2.4), (S2.5), and (S2.6), with $\beta_1 = 0$ and $\beta_2 = 1$. Note that with this choice for $\beta_1$ and $\beta_2$, the Beta function and the incomplete Beta function in equation (S2.4) are not well defined separately, but the integral that they represent when combined does exist (similar for equation (S2.6) if $n_{aB} = 0$). Furthermore, note that the incomplete Beta function in equation (S2.5) does not exist if $n_{aB} = 0$. This is in line with the use of the non-recurrent mutation model in the literature. Since we designate $b$ to be the ancestral allele at the neutral locus, a configuration with $n_{aB} = 0$ corresponds to a configurations with all alleles at the neutral locus being ancestral. In the non-recurrent mutation model there is an infinite supply of ancestral allele, and thus the expected number of such configurations is infinite. To arrive at a proper probability distribution over all two-locus haplotype configurations, we employ the following strategy. We first compute the likelihoods for all configurations with $n_{aB} > 0$, and define $m$ as 1 minus the sum of all these likelihoods. We than distribute the mass $m$ among the remaining configurations with $n_a = 0$ (and $n_{AB} = 0$) proportional to the probability of sampling $n_{Ab}$ alleles of type $A$ given $x_A$.

###### S2.4.4 Non-Stationary Distribution

In the case when $p_B(x)$ is a non-stationary distribution, for example considering a beneficial mutation which is introduced at time $\eta_3$ in Figure 2 in the main text, directly computing $p_B(x)$ is in general impractical. However, to approximate $p_B(x)$ we assume that $p_B(x)$ follows a Beta$(\beta_1, \beta_2)$ distribution and estimate $\beta_1$ and $\beta_2$ using the method of moments. Namely, we consider the moments of orders $n = 1$ and $n = 2$ for $p_B(x)$ and select $\beta_1$ and $\beta_2$ such that a Beta$(\beta_1, \beta_2)$ distribution has the same moments. This is then used to derive the initial moments using the method outlined in Section S2.4.2. We find that this approximation works well in practice.

###### S2.4.5 Two-Locus Stationary Distribution

In addition to modeling the setting in which a selected allele arises on a background which is at stationarity, one may be interested in modeling a situation in which an existing neutral allele gains a selective advantage/disadvantage and selection starts from standing variation. In such a setting, the ODEs need to be initialized from the two-locus stationary distribution. To do this, one can take a similar approach as in `LDpop` (Kamm et al., 2016). Namely, one can formulate a two-locus Moran model which accounts for genetic drift, mutation, and recombination and compute its transition matrix. This transition matrix can then be used to compute the stationary distribution of the moments by either solving the appropriate set of linear equations or via power iteration, which can be more efficient for sparse matrices. For an exact result, one would use the augmented Moran model of Kamm et al. (2016) which includes partially specified haplotypes, while for larger sample sizes ($n \geq 20$) the normal two-locus Moran model of Ethier and Kurtz (1993) serves as a good approximation.

### S3 Additional Simulation Results

In this section we display some additional results for some parameter configurations not shown in the main text.

#### *S3.1 Fixed Initial Frequencies*

###### S3.1.1 Additional Parameter Combinations

In Figure S1, we present the expected frequency of haplotype $AB$ with the same parameters as Figure 5 in the main text but $\rho = 0.0004$. We can see that since the two loci are tightly linked the trajectories closely resemble expected trajectory of the $A$ allele in Figure 4 in the main text.

###### S3.1.2 Comparing to Expected Trajectories under Exact Wright-Fisher Diffusion

To further validate the output from the numerical solutions to the ODEs, we compared them to the expected allele frequency trajectories of the one-locus Wright-Fisher diffusion, computed with high accuracy. We obtained these trajectories using an implementation of the spectral method developed by Steinrücken et al. (2014). This method can be used to compute the expected trajectories in the one-locus case with high accuracy for constant population size history. This can be achieved by computing the likelihood of a temporal sample of size one, with one copy of the derived allele, since this corresponds to the binomial integral that also yields the expected allele frequency. In Figure S2, we compare the expected allele frequencies of a selected allele calculated from the moment ODE, simulations using `SimuPOP`, and the exact Wright-Fisher diffusion for a population of size 10,000 over 6,000 generations across three different selection parameters, with initial frequency 0.05. This figure aligns with the first 6,000 generation of the demography shown in Figure 2 and used in Figure 4 in the main text. The solutions from the moment ODE and the Wright-Fisher diffusion show a high level of agreement. This indicates that, in the respective parameter range, the discrepancies between the moment ODE and `SimuPOP` are only due to noise from random sampling and the fact that the Wright-Fisher diffusion is only an approximation of the finite Wright-Fisher model. Since the ODEs are derived based on the diffusion, they can only be as good an approximation as the diffusion is. These

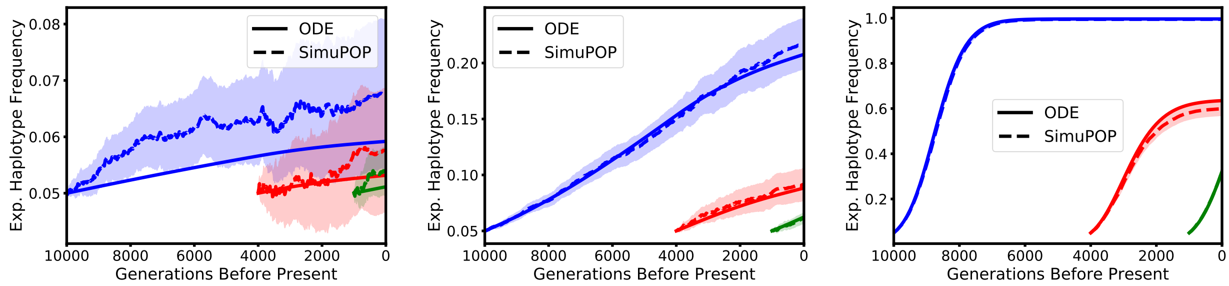


(a) Selection coefficient $\sigma = 1$. (b) Selection coefficient $\sigma = 10$. (c) Selection coefficient $\sigma = 100$.

Figure S1: Expected frequency of the $AB$ haplotype, but for strong linkage ($\rho = 0.0004$). Again, blue, red, and green correspond to the dynamics starting 10,000 generations, 4,000 generations (beginning of the bottleneck), and 1,000 generations before present (beginning of exponential growth), respectively. The dotted and solid lines correspond to the simulation average and ODE solution, respectively, with the shaded region indicating the 95% confidence interval.

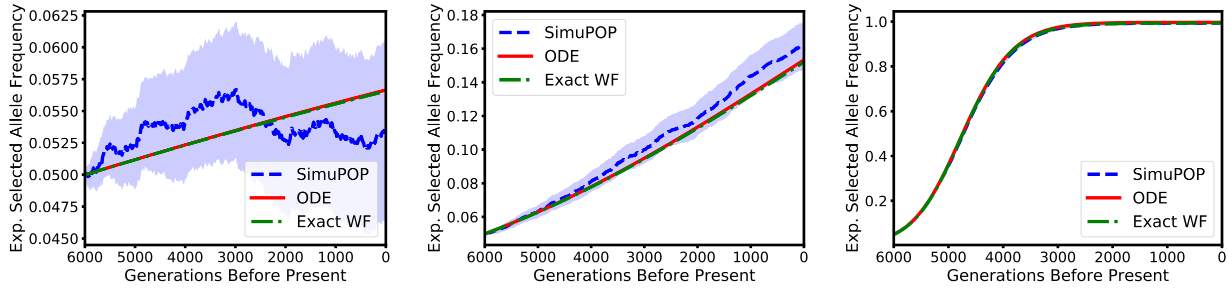


(a) Selection coefficient $\sigma = 1$. (b) Selection coefficient $\sigma = 10$. (c) Selection coefficient $\sigma = 100$.

Figure S2: Expected allele frequencies selected allele calculated from numerical solutions to the moment ODE (blue dashed line), simulations using `SimPOP` (red solid line), and exact Wright-Fisher diffusion (green dash-doted line) for a population of size 10,000 over 6,000 generations for three different selection parameters. Shaded areas indicate 95% confidence.

conclusions are also supported by the results presented in Figure S3, which displays the expected trajectories of the selected allele derived from the moment ODE, `SimuPOP`, and the exact Wright-Fisher diffusion, but for a population of size 2,000 over 3,000 generations, aligning with the first 3,000 generations of the red trajectories in Figure 4 in the main text.

##### *S3.2 Neutral Locus at Stationarity*

In this section we present analogous results to those displayed in Section 3.1.2 in the main text, with the selected allele $A$ is initialized at frequency 0.03 and 0.01 instead of 0.05. Figures S4 and S5 display the heterozygosity over time (analogous to Figure 8 in the main text) with $A$ initialized from 0.03 and 0.05, respectively. Similarly, Figures S6 and S7 display $D^2$ over time (analogous to Figure 9 in the main text) where $A$ is initialized at frequency 0.01 and 0.03, respectively.

##### *S3.3 Site-Frequency-Spectrum in a Genomic Window*

Here we present additional figures to supplement the discussion in Section 3.2 in the main text. Figure S8 shows the folded SFSs in the scenarios exhibited in Figure 10 in the main text, computed using the recurrent

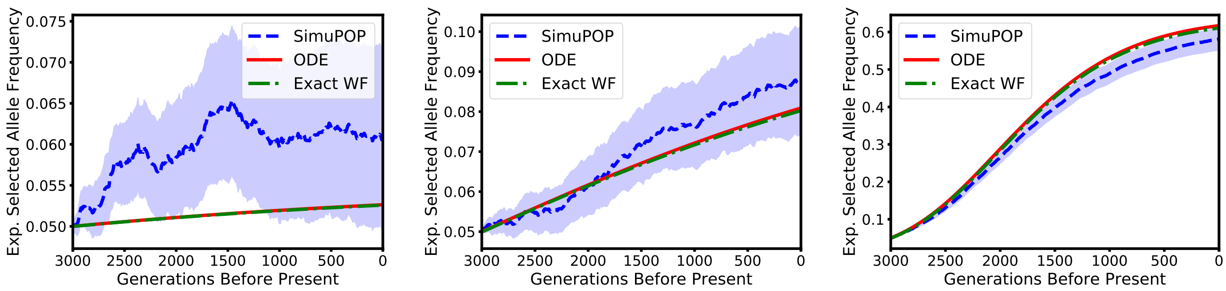


(a) Selection coefficient $\sigma = 1$.  (b) Selection coefficient $\sigma = 10$.  (c) Selection coefficient $\sigma = 100$.

Figure S3: Expected allele frequencies selected allele calculated from numerical solutions to the moment ODE (blue dashed line), simulations using `SimPOP` (red solid line), and exact Wright-Fisher diffusion (green dash-doted line) for a population of size 2,000 over 3,000 generations for three different selection parameters. Shaded areas indicate 95% confidence.

mutation model. Each entry of the SFS is given by

$$S_k^n(t) := \sum_{\substack{i \in E \\ i_B = k}} M_i(t) + \sum_{\substack{i \in E \\ i_b = k}} M_i(t).$$

### S4 Approximating Moments for Large Difference of Orders

We applied the regular *logit-linear* moment approximation to estimate moments where the difference in order is large, for example, to approximate moments of order 101 from order 31. We observed a loss in accuracy, since too much probability mass is shifted away from the boundaries of the simplex, that is, configurations where at least one of the possible haplotypes is not present in the sample. To remedy these inaccuracies, we have developed a prototype approximation method for higher-order moments. In this approach, we first use the regular *logit-linear* for the interior and the boundaries separately, where the different boundaries are classified by the number of haplotypic classes are unobserved in the respective sample configuration. Note that the interior is the "boundary" where all haplotypic classes are observed. We then combine these separate approximations into one approximation for the entire vector of moments as follows. Starting from the interior and adding different boundaries successively adds additional entries into the vector on each level. The probability mass for these new entries is always taken out of the boundary on the next level. Proceeding iteratively, we arrive at the final vector with entries for all configurations. This procedure does shift probability mass from the boundaries into the interior, but less than the regular *logit-linear* would have, thus resulting in a better approximation. In Figure S9 we present the results of applying this technique to approximate SFSs for a sample of size 101, computed from a moment vector of order 31, and assess the accuracy by comparing to simulations using `SLiM`, see Section 3.2 in the main text for a full description. We omit the case $\sigma = 100$, because we did observe that the new approximation became numerically unstable for large $\sigma$, which we will address in future work. The SFS obtained from the numerical solution of the ODE combined with the new moment approximation (dotted lines) are generally in good agreement with the simulated SFS (solid lines).

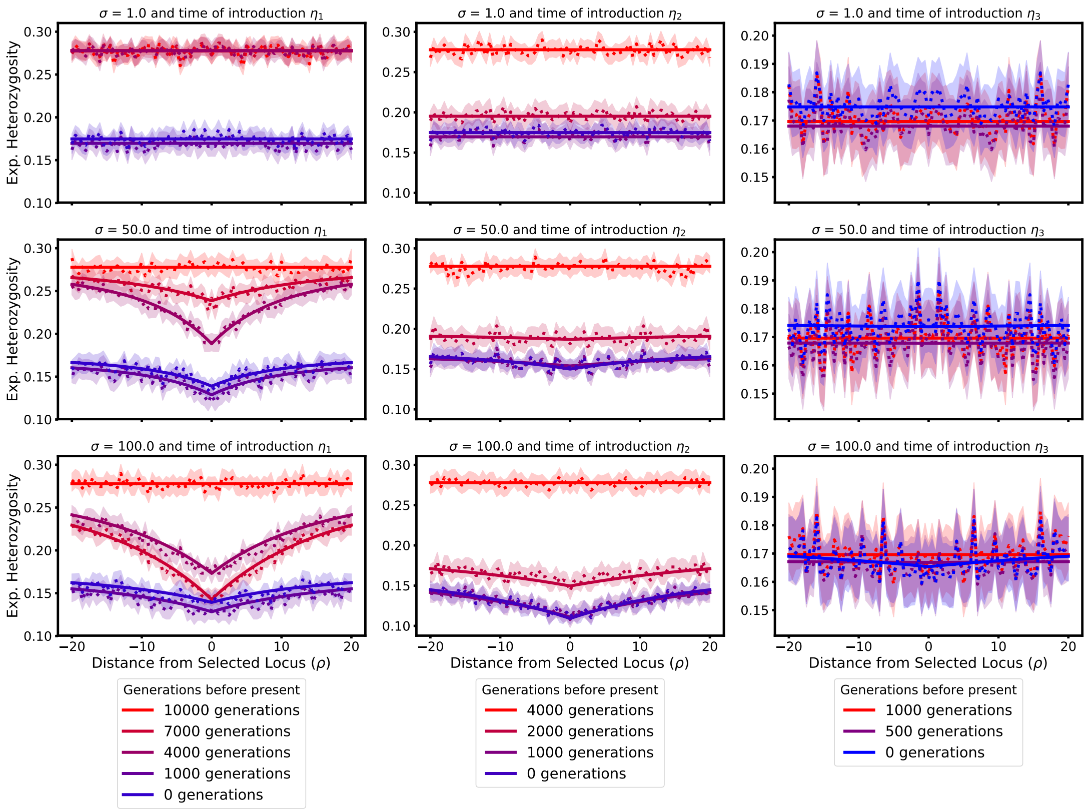


Figure S4: Expected heterozygosity across a 100 kbp region with different selection coefficients and starting times. The beneficial allele is introduced at frequency $x_A = 0.03$. The dotted and solid lines represent the simulation and ODE results, respectively, with the shaded region representing a 95% confidence interval.


Kamm, J. A., Spence, J. P., Chan, J., and Song, Y. S. (2016). Two-locus likelihoods under variable population size and fine-scale recombination rate estimation. *Genetics*, **203**(3), 1381–1399.

Kincaid, D., Kincaid, D. R., and Cheney, E. W. (2009). *Numerical Analysis: Mathematics of Scientific Computing*, volume 2. American Mathematical Society.

Klenke, A. (2008). *Probability Theory : A Comprehensive Course*. Springer, London.

Steinrücken, M., Bhaskar, A., and Song, Y. S. (2014). A novel spectral method for inferring general diploid selection from time series genetic data. *Annals of Applied Statistics*, **8**(4), 2203–2222.

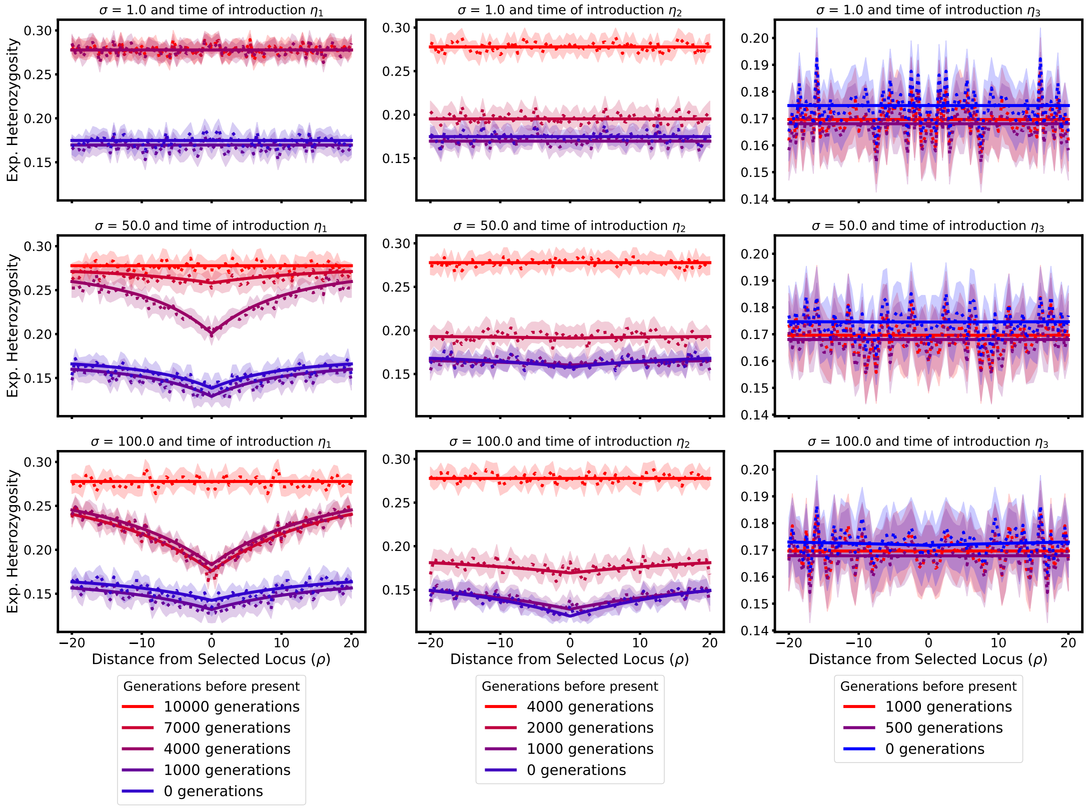


Figure S5: Expected heterozygosity across a 100 kbp region with different selection coefficients and starting times. The beneficial allele is introduced at frequency $x_A = 0.01$. The dotted and solid lines represent the simulation and ODE results, respectively, with the shaded region representing a 95% confidence interval.

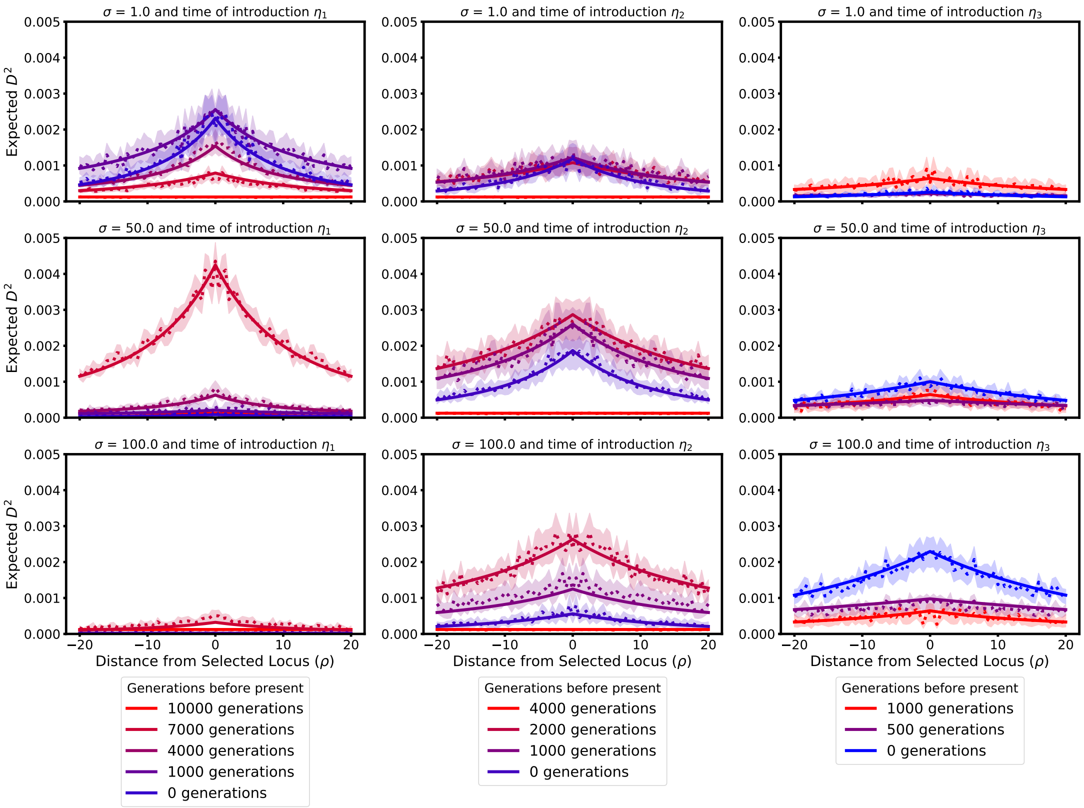


Figure S6: $D^2$ across a 100 kbp region with different selection coefficients and starting times. The beneficial allele is introduced at frequency $x_A = 0.03$. The dotted and solid lines represent the simulation and ODE results, respectively, with the shaded region representing a 95% confidence interval.

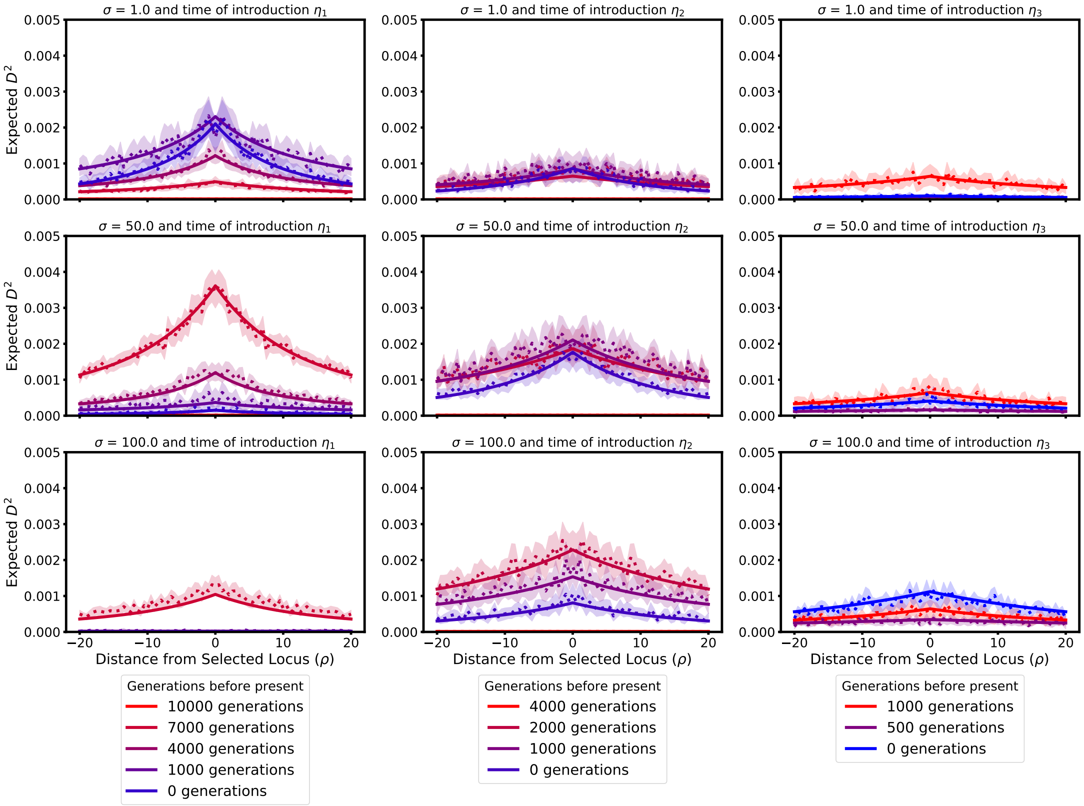


Figure S7: $D^2$ across a 100 kbp region with different selection coefficients and starting times. The beneficial allele is introduced at frequency $x_A = 0.01$. The dotted and solid lines represent the simulation and ODE results, respectively, with the shaded region representing a 95% confidence interval.

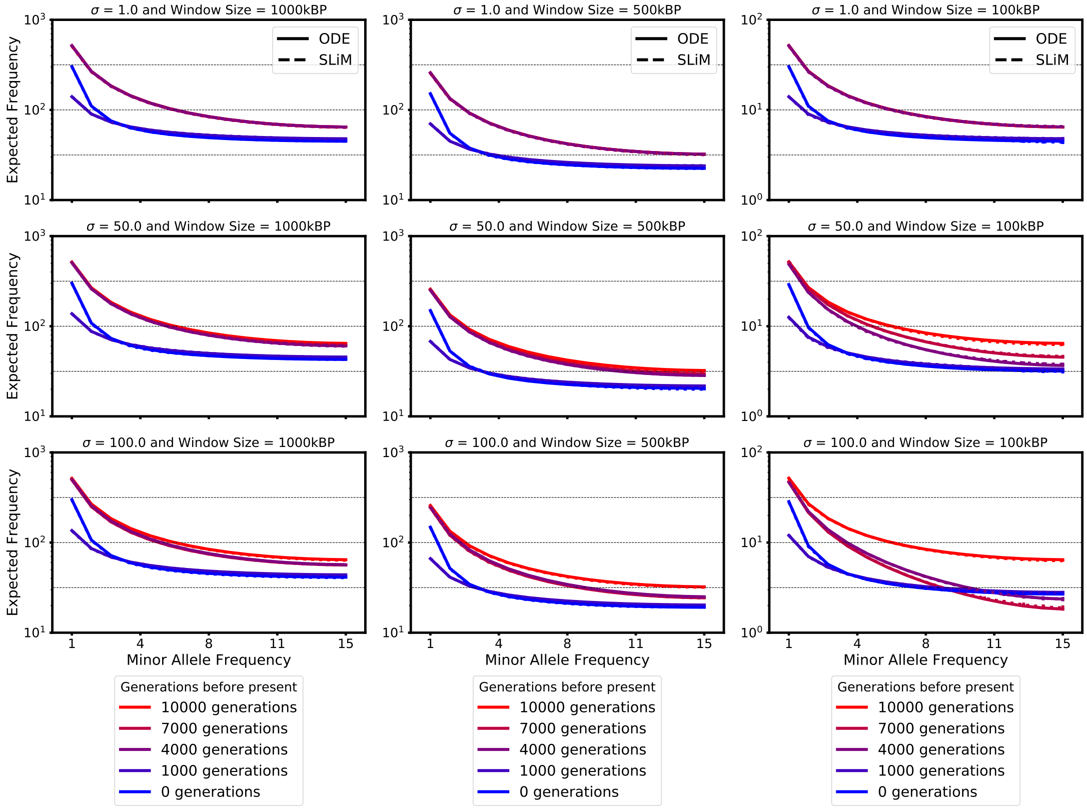


Figure S8: Local folded site-frequency-spectra for windows of size 100 kbp, 500 kbp, and 1 Mbp, for selection coefficients $\sigma = 1$, 50, and 100. Computed using the recurrent mutation model. The demographic history is given in Figure 2 in the main text and the beneficial allele is introduced 10,000 generations before present ($\eta_1$).

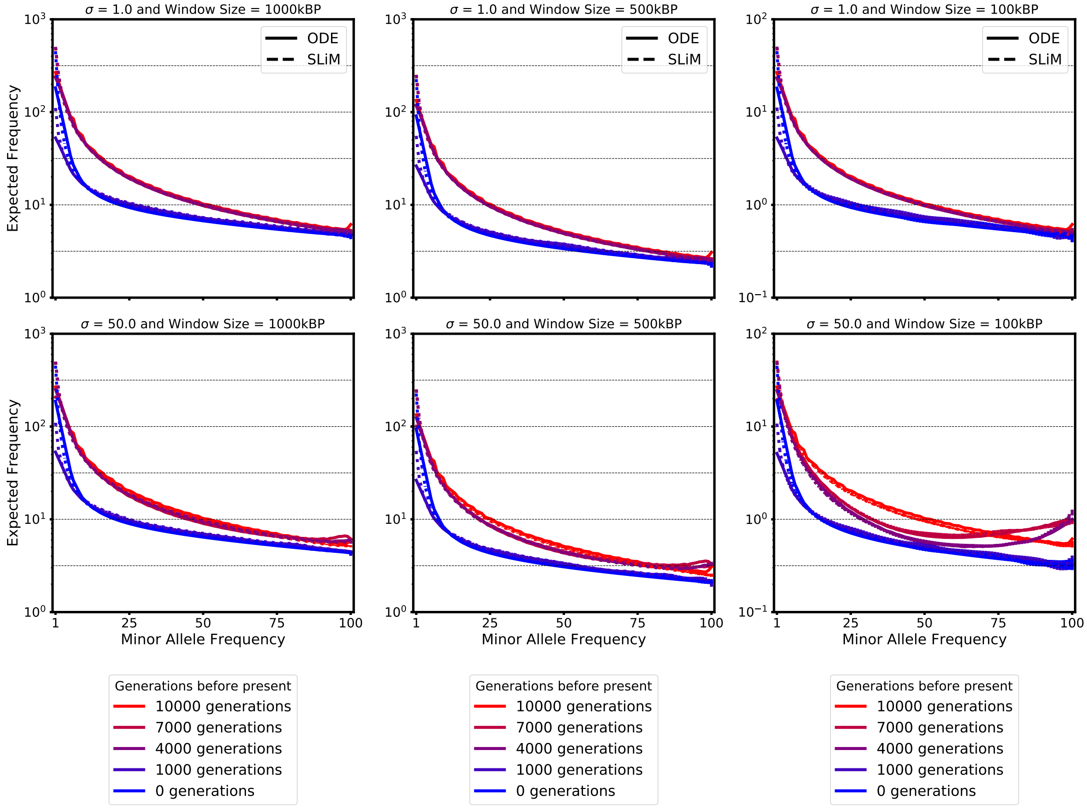


Figure S9: Local site-frequency-spectra for windows of size 100 kbp, 500 kbp, and 1 Mbp, for sample size 51, and for selection coefficients $\sigma = 1$ and 50. The demographic history is given in Figure 2 in the main text and the beneficial allele is introduced 10,000 generations before present ($\eta_1$). Note that for the third columns the y-axis limits have changed.
